# Subset-specific mitochondrial and DNA damage shapes T cell responses to fever and inflammation

**DOI:** 10.1101/2022.11.14.516478

**Authors:** Darren R. Heintzman, Joel Elasy, Channing Chi, Xiang Ye, Evan S. Krystoviak, Wasay Khan, Lana Olson, Angela Jones, Kelsey Voss, Andrew R. Patterson, Ayaka Sugiura, Frank M. Mason, Hanna S. Hong, Lindsay Bass, Katherine L. Beier, Wentao Deng, Costas A. Lyssiotis, Alexander G. Bick, W. Kimryn Rathmell, Jeffrey C. Rathmell

## Abstract

Heat is a cardinal feature of inflammation. Despite temperature variability and dependence of enzymes and complexes, how heat and fever affect immune cells remains uncertain. We found that heat broadly increased inflammatory activity of CD4^+^ T cell subsets and decreased Treg suppressive function. Th1 cells, however, also selectively developed mitochondrial dysfunction with high levels of ROS production and DNA damage. This led Th1 cells to undergo *Tp53*-dependent death, which was required to minimize the accumulation of mutations in heat and inflammation. Th1 cells with similar DNA damage signatures were also detected in Crohn’s disease and rheumatoid arthritis. Fever and inflammation-associated heat thus selectively induce mitochondrial stress and DNA damage in activated Th1 cells that requires p53 to maintain genomic integrity of the T cell repertoire.

**One Sentence Summary:** Fever temperatures augment CD4^+^ T cell-mediated inflammation but induce differential metabolic stress and DNA damage in T cell subsets, with Th1 cells selectively sensitive and dependent on p53 to induce apoptosis and maintain genomic integrity.

## Introduction

Cell metabolism directly shapes T cell differentiation and function, with each T cell subset requiring a specific metabolic program (*1*). These processes are underpinned by wide ranging enzymatic reactions and protein complexes evolved to operate optimally within narrow biophysical and structural windows, including substrate or co-factor availability, redox, pH, and temperature (*2, 3*). However, temperature fluctuations occur frequently, particularly during instances of immune challenge. In addition to systemic fevers, locally inflamed tissues have long been recognized to establish and maintain elevated temperatures as a cardinal sign of inflammation (*4*). This is particularly well documented in chronic autoimmune diseases such as rheumatoid arthritis (*5, 6*). Even under normal circumstances, temperature is not consistent over time or across body locations and can fluctuate dramatically in extremities and in response to environmental conditions (*7*). While T cells are programmed to adapt to increasing temperatures through induction of a protective heatshock response (*8*) that may be induced in T cells at lower temperatures than other immune cells (*9*), the effect of heat and febrile temperatures on T cell metabolism and subsets remains poorly understood.

Fever temperatures may impact T cells through altered signaling, stress, or metabolic pathways. Indeed, recent studies have suggested increased Th2 T cell differentiation and function at febrile temperatures (*10*). Heat-exposed CD4^+^ Th17 cells were also shown to selectively increase differentiation and pathogenicity following altered SMAD signaling (*11*). T cells are reported to have increased migration in fever conditions through HSP90-α4 integrin interactions and increased adhesion (*12–16*), and transient heat can enhance CD8^+^ T cell function through increased mitochondrial protein translation and respiratory capacity (*17*). The distinct metabolic programs of each T cell subset and broad dependence of those programs on temperature-sensitive enzymes now suggests that T cells will be subject to subset-specific responses to heat-induced metabolic adaptations.

Here we tested the hypothesis that febrile temperatures selectively alter the metabolism and fate of CD4^+^ T cell subsets. While heat broadly increased T cell-mediated inflammatory capacity and proliferation, Th1, Th17, and Treg subsets each exhibited distinct metabolic adaptations and Th1 cells selectively experienced high levels of mitochondrial stress and ROS. This led to double stranded DNA breaks that required the activation of p53 to eliminate damaged cells and prevent the accumulation of excessive mutations. Mutagenesis thus appears to be a common feature of heat in Th1 cells that renders p53 critical to maintain genomic integrity of the T cell repertoire and reduce risk for transformation in inflammation.

## Results

### CD4^+^ T cell functions favor inflammatory states at febrile temperatures

To establish the effect of heat on T cell function, CD4^+^ T cells were cultured at either 37°C or 39°C in activating (Th0) or subset differentiating (Th1, Th17, and induced Treg, iTreg) conditions. Cells were cultured for 3-4 days to allow sufficient time for subset differentiation. Interestingly, we observed a greater percentage of Th1 and Th0 cells producing IFNγ^+^ at 39°C (Figure 1A, B). As previously reported (*11*), Th17 cells expressed significantly more IL17a when cultured at 39°C compared to 37°C and no change was observed in expression of FOXP3 in iTregs (Figure 1C, D). Recently published work has suggested that regulatory T cells are more stable *ex-vivo* when cultured at lower temperatures (*18*). While FOXP3 expression was unchanged with heat, iTregs cultured at 39°C were less effective at suppressing effector CD8^+^ T cell proliferation (Figure 1E). All T cell subsets had greater rates of proliferation at 39°C as determined by dilution of Cell Trace Violet (CTV) and increased division index (Figure 1F, G). Consistent with a need for T cell activity in inflammatory settings, effector functions are thus broadly increased while suppressive functions are decreased in CD4^+^ T cell exposed to elevated temperatures.

**Figure 1:**
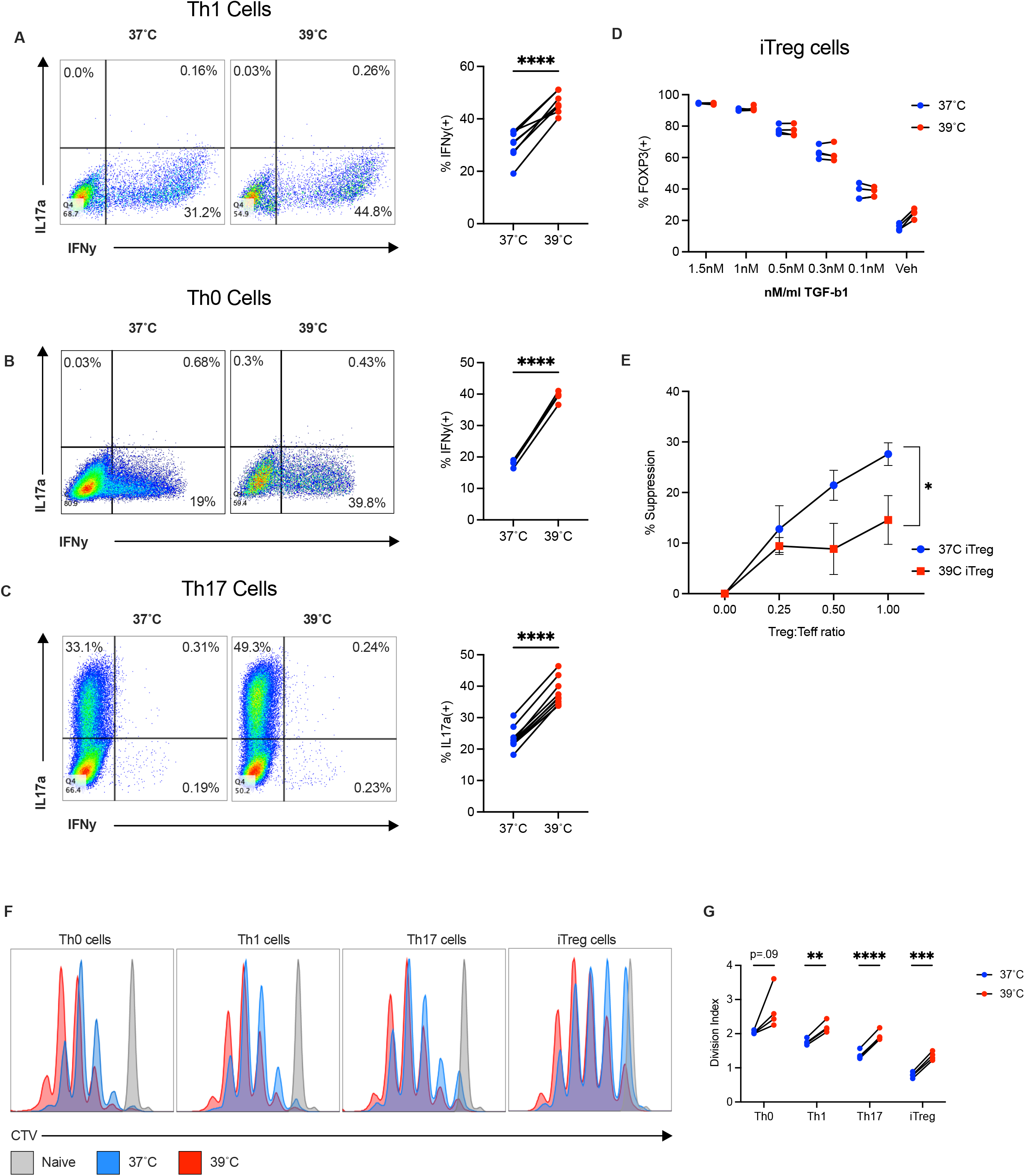
CD4^+^ T cell function is broadly altered at febrile temperatures. A) IL17a expression by Th17 cells at 37°C and 39°C B) FOXP3 expression of iTreg cells cultured in dilution series of hTGF-β (C) Suppression assay of CD8^+^ T cells co-cultured with CD4^+^ iTregs cultured at 37°C or 39°C. D) IFNγ expression by Th1 cells at 37°C and 39°C E) IFNγ expression by Th0 cells at 37°C and 39°C. F) Cell Trace Violet staining of CD4^+^ T cell subsets cultured at 37°C (Blue) or 39°C (Red). Naïve (Grey) serves as nonproliferating control G) Division index calculated using CTV data in (F). Biological replicates shown with standard deviation and mean (*, P < 0.05; **, P < 0.01; ***, P < 0.001, ****, P < 0.0001; A-C, G: Paired T-Test, E: Multiple unpaired T-Tests)

### Heat selectively enhances glycolysis in Th17 and iTreg cells

Given links between metabolism and T cell function and temperature dependence of enzymes, we hypothesized that higher temperatures would alter cellular metabolism to influence cell fate. The Akt/mTORC1 pathway plays key roles in T cell function and metabolism and thus was assessed. The phosphorylation and activation of mTOR pathway components including Akt, S6, and 4EBP1 were elevated in all CD4^+^ T cell subsets (Figure 2A, S1A-C). To broadly measure glucose utilization, expression of the glucose transporter Glut1 was assayed by flow cytometry. Interestingly, Glut1 expression was only significantly enhanced in Th17 and iTreg subsets when cultured at 39°C (Figure 2B). We examined glycolytic rates by measuring extracellular flux and acidification rates (ECAR). Consistent with increased Glut1 expression, ECAR was selectively increased in Th17 and iTreg cells at 39°C (Figure 2C-E). These data are consistent with recent findings of CD8^+^ T cells, where increased glycolysis was measured in cells cultured at febrile temperatures (*17*). In contrast to Th17 and iTreg cells, Glut1 expression and ECAR of Th1 cells were largely unchanged by heat. This subset-specific increase in glycolysis was corroborated when lactate secretion by Th17 and iTreg cells was measured directly (Figure S1D). While a modest increase in oxygen consumption rates (OCR) was observed in iTreg cells at febrile temperatures, Th1 cells showed a modest decrease in maximal OCR and overall changes in OCR were minimal (Figure S1E, F). These data show that CD4^+^ T cells metabolically adapt to increased temperatures in subset-specific manners, with Th17 and iTreg cells becoming significantly more glycolytic while glycolysis and respiration remain largely unchanged in Th1 cells.

**Figure 2:**
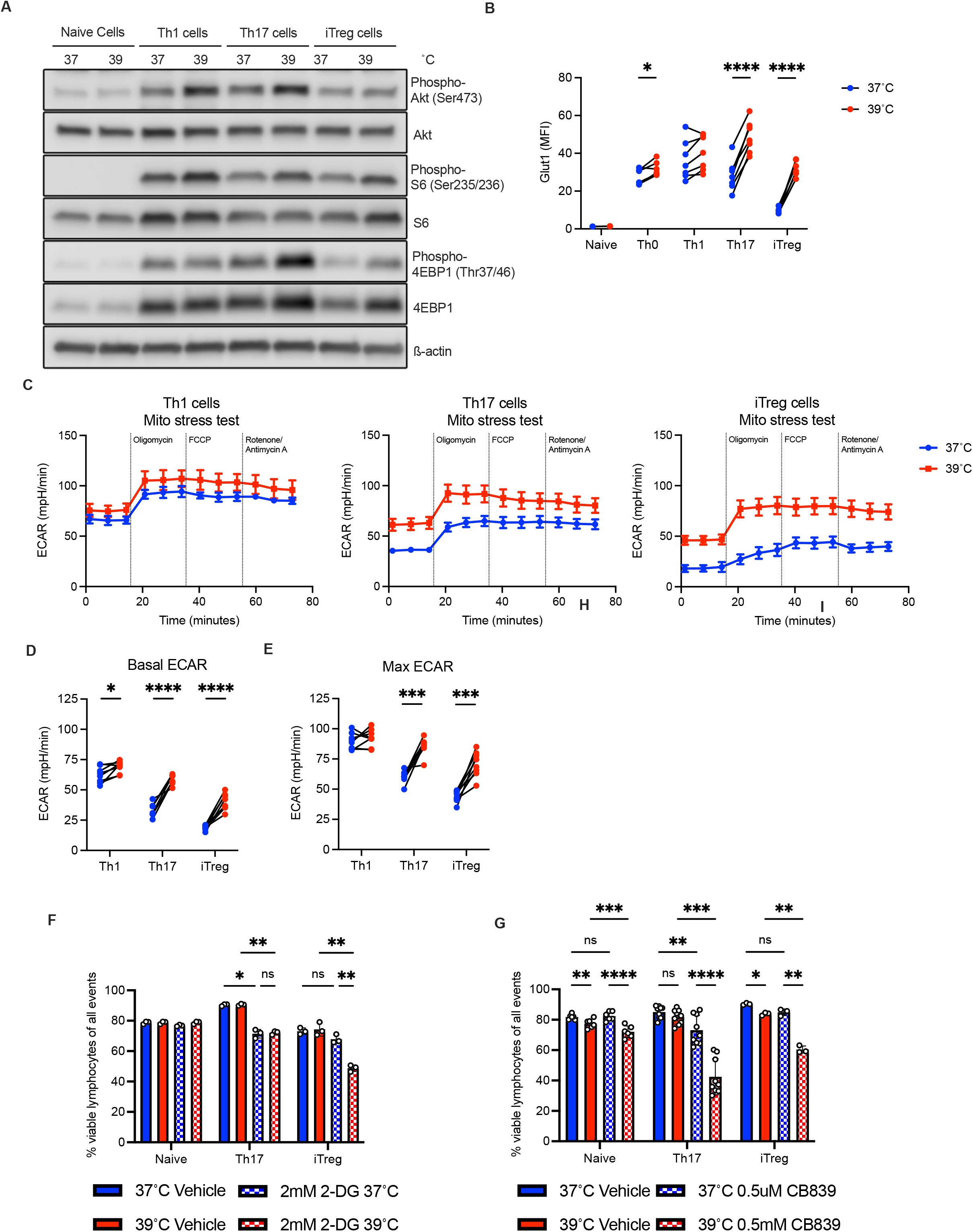
CD4^+^ T cells rely on altered, differential metabolic programming at febrile temperatures. A) Phosphorylation of components of mTOR pathway, assessed by western blotting. B) Glut1 expression measured by flow cytometry C) Representative bioreplicates of Th1, Th17, and iTreg ECAR measured by Seahorse. D, E) Basal and maximal ECAR measured by Seahorse F) Viability of naïve, Th17, and iTreg cells with or without 2-DG in culture at 37 or 39°C G) Viability of Naïve, Th17 and iTreg cells cultured with or without CB-839. Biological replicates shown with standard deviation and mean (*, P < 0.05; **, P < 0.01; ***, P < 0.001, ****, P < 0.0001; B, D, E, H: Paired T-Test, F, G: 2-Way ANOVA)

### Th17 and iTreg cells rely on differential metabolic programs at febrile temperatures

To better understand heat-induced metabolic adaptations, tricarboxylic acid cycle and related metabolites were measured in naïve, Th17, and iTreg cells cultured at 37°C and 39°C. Glutaminolysis and TCA cycle intermediates were significantly enriched in Th17 cells at 39°C compared to culture at 37°C, while iTregs showed modest increases in these pathways (Figure S1G). Th17 and iTreg cells were cultured for 3 days at 37°C or 39°C with metabolic inhibitors to test the dependence of Th17 and Treg on glycolysis and glutaminolysis. Cells were first treated with the non-hydrolyzable glucose analog 2-deoxyglucose (2-DG). 2-DG mildly decreased FOXP3 induction in iTreg, but augmented IL17a production of Th17 cells at both 37°C and 39°C (Figure S1H). 2-DG also decreased the viability of iTregs at 39°C, while Th17 were unaffected (Figure 2F). Because Th17 cells can efficiently use glutamine as a fuel (*19*), we hypothesized that Th17 cells may instead rely on this pathway at 39°C. To test this, cells were cultured with the Glutaminase 1 (Gls) inhibitor, CB839, for 3 days and cytokine production and viability were assessed. Th17 cytokine production was inhibited with the addition of CB839, and this remained the case at 39°C (Figure S1I). While CB839 only slightly affected cell viability at 37°C, Th17 cells showed a significant loss of viability at 39°C with Gls inhibition (Figure 2G). Treg, which do not respond to Gls inhibition at 37°C, were also affected, although to a lesser extent. These data show that Th17 and iTreg cells broadly alter metabolic pathway reliance at 39°C, with iTreg increasingly dependent on glycolysis and Th17 more dependent on glutaminolysis.

### Heat selectively induces mitochondrial dysfunction in Th1 cells

Because CD8^+^ T cells increase mitochondrial mass when exposure to heat (*17*), we next examined CD4^+^ T cell mitochondria content and function. Consistent with this previous study, all activated CD4^+^ T cell subsets had increased overall mitochondrial mass at 39°C when assessed by flow cytometry and the probe Mitotracker green (Figure S2A). Electron microscopic imaging revealed that mitochondria in Th0 and Th1 subsets developed greater heterogeneity in mitochondrial density and cristae formation, indicative of cells accumulating damaged or dysfunctional mitochondria (Figure 3A, B, S2B). Quantification confirmed that larger mitochondria accumulated in many Th1 cells cultured at 39°C (Figure 3A, B). Th17 cells also exhibited mitochondrial morphological changes at 39°C, while iTreg mitochondria appeared unchanged at 39°C (Figure S2B). Mitochondrial superoxide is a marker of mitochondrial dysfunction and is generated by dysfunctional or inefficient electron transport (*20*). When measured using the fluorescent probe MitoSOX, mitochondrial superoxide was sharply and selectively elevated in only Th0 and Th1 subsets (Figure 3C).

**Figure 3:**
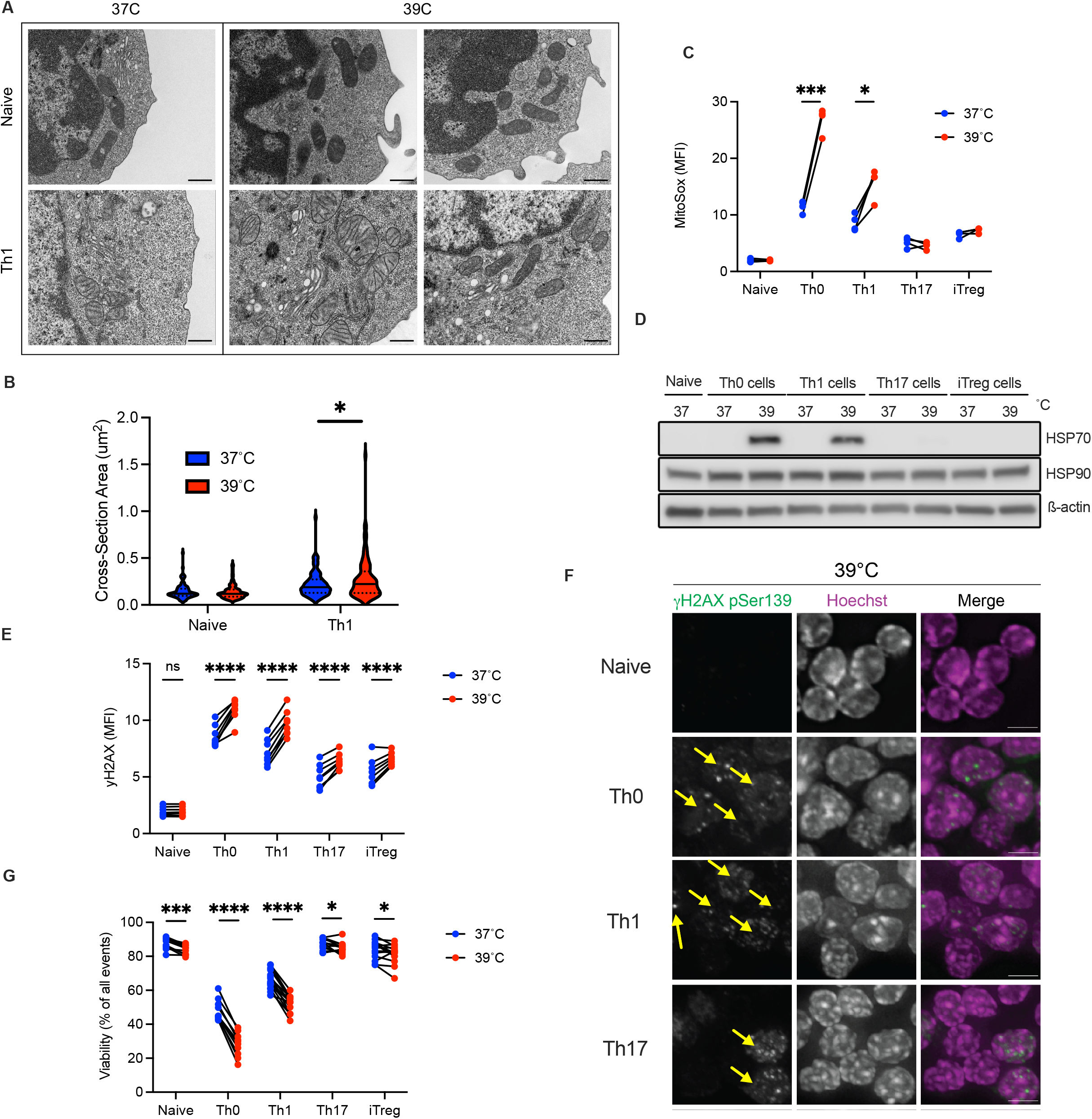
Th1 mitochondrial dysfunction leads to differential response to heat stress. A) Electron microscopy imaging of naïve and Th1 mitochondria at 37°C and 39°C. B) Mitochondrial area quantified from electron microscopic imaging C) Mitochondrial superoxide measured by MitoSOX red D) HSP70, HSP90, and b-Actin expression assessed by western blot. E) DNA damage was measured by γH2AX staining via flow cytometry F) γH2AX foci assessed by confocal microscopic imaging in cells cultured at 39°C. G) Viability was measured using live-dead exclusion dye by flow cytometry. Quantified by live cells as a percentage of total events. Scale bars are 500 nm for (A), 5um for (F). Biological replicates shown with standard deviation and mean (*, P < 0.05; **, P < 0.01; ***, P < 0.001, ****, P < 0.0001; Paired T-Test)

Superoxide production in Th0 and Th1 subsets cultured at 39°C could lead to a stress response or cause DNA damage. To measure stress responses, the heat shock proteins HSP70 and HSP90 were assessed through flow cytometry and western blotting. While HSP90 was constitutively expressed in all cell types at both temperatures, HSP70 was selectively induced in Th0 and Th1 cells cultured at 39°C (Figure 3D, S3A, B). Reactive oxygen species can cause DNA damage and double stranded breaks leading to the phosphorylation of H2AX to form γH2AX (*21*). Indeed, Th0 and Th1 cells displayed higher γH2AX staining at both 37°C and 39°C that did not occur in Th17 or Treg (Figure 3E). While heterogeneous at cell populations levels, Th1 and Th0 cells had notably higher frequencies of γH2AX^+^ cells at febrile temperatures than other subsets (Figure 3E, F, S3C). These data are consistent with increased ROS and DNA damage that have been observed in T cells in diseases associated with inflammation and resulting temperature change, including rheumatoid arthritis (*22–24*).

Consistent with increased markers of stress and DNA damage, viability of Th0 and Th1 subsets decreased significantly at 39°C, while naïve, Th17, and iTreg subset viability remained high (Figure 3G). Viability in Th0 and Th1 subsets was temperature dependent, as viability decreased in a stepwise manner as culture temperatures increased from 30°C to 39°C (Figure S4A). Loss of cell viability was not dependent on accumulation of IFNγ in cell cultures, as high levels of neutralizing anti-IFNγ antibody had no effect on viability in the most severely affected Th0 condition (Figure S4B). Interestingly, while iTreg viability at 39°C was TGF-β dependent as T cells failed to differentiate and upregulate FoxP3 in the absence of TGF-β (Figure S4C), removal of TGF-β from culture did not meaningfully impact Th17 viability at 39°C (Figure S4D). IL17a production was, however, strongly inhibited by loss of TGF-β (Figure S4E). These data highlight that T cells adapt to temperature change in subset specific ways, with Th0 and Th1 cells accumulating significantly more cell stress and resultant loss of viability.

### Mitochondrial ROS induced DNA damage activates p53 in Th1 cells

To determine the molecular mechanism of cell death, we performed an *in vitro* CRISPR Screen in Th1 cells using a custom-guide RNA library focused on 53 cell death-related genes, including apoptosis, necroptosis, and ferroptosis pathways. T cells were activated and transduced with the cell death gRNA library at 39°C and guide frequencies were measured by DNA sequencing to determine if disruption of a specific gene altered the fitness of T cells at fever temperatures. Surprisingly, the p53 pathway was highly enriched, with deletion of *Trp53, Puma* (*Bbc3*), *Bax* (*Bcl2l11*), and *Fas*, all providing a survival or proliferative advantage for Th1 cells at 39°C (Figure 4A). p53 is activated by ATM or ATR-mediated phosphorylation and analysis of p53 by immunoblotting revealed phosphorylation of serine-15 on p53 was significantly increased in Th0 and Th1 cells at 39°C while total p53 remained unchanged (Figure 4B, S5A, B). In contrast, Th17 and iTreg cells were unaffected at either temperature. Expression of the p53 target, p21, was also significantly enhanced at 39°C selectively in Th0 and Th1 cells (Figure 4B, Figure S5C), supporting selectively enhanced p53 transcriptional activity in these subsets. To validate the results of the CRISPR screen, T cells were isolated from age-matched wild-type and Tp53^-/-^ mice and cultured at 37°C and 39°C to assess viability. Indeed, p53-deficiency partially restored viability in Th0 and Th1 cells at 39°C, while viability of naïve, Th17, and iTreg cells was unaffected by p53-deficiency at either temperature (Figure 4C). Although viability was partially rescued in Tp53^-/-^ cells, Th0 and Th1 subsets still displayed the same, or even enhanced, markers of cell stress at 39°C compared to wild type including increased ROS and γH2AX (Figure S6A, B). Additionally, Tp53^-/-^ cells had greater effector cytokine expression at 39°C (Figure S6C-H).

**Figure 4:**
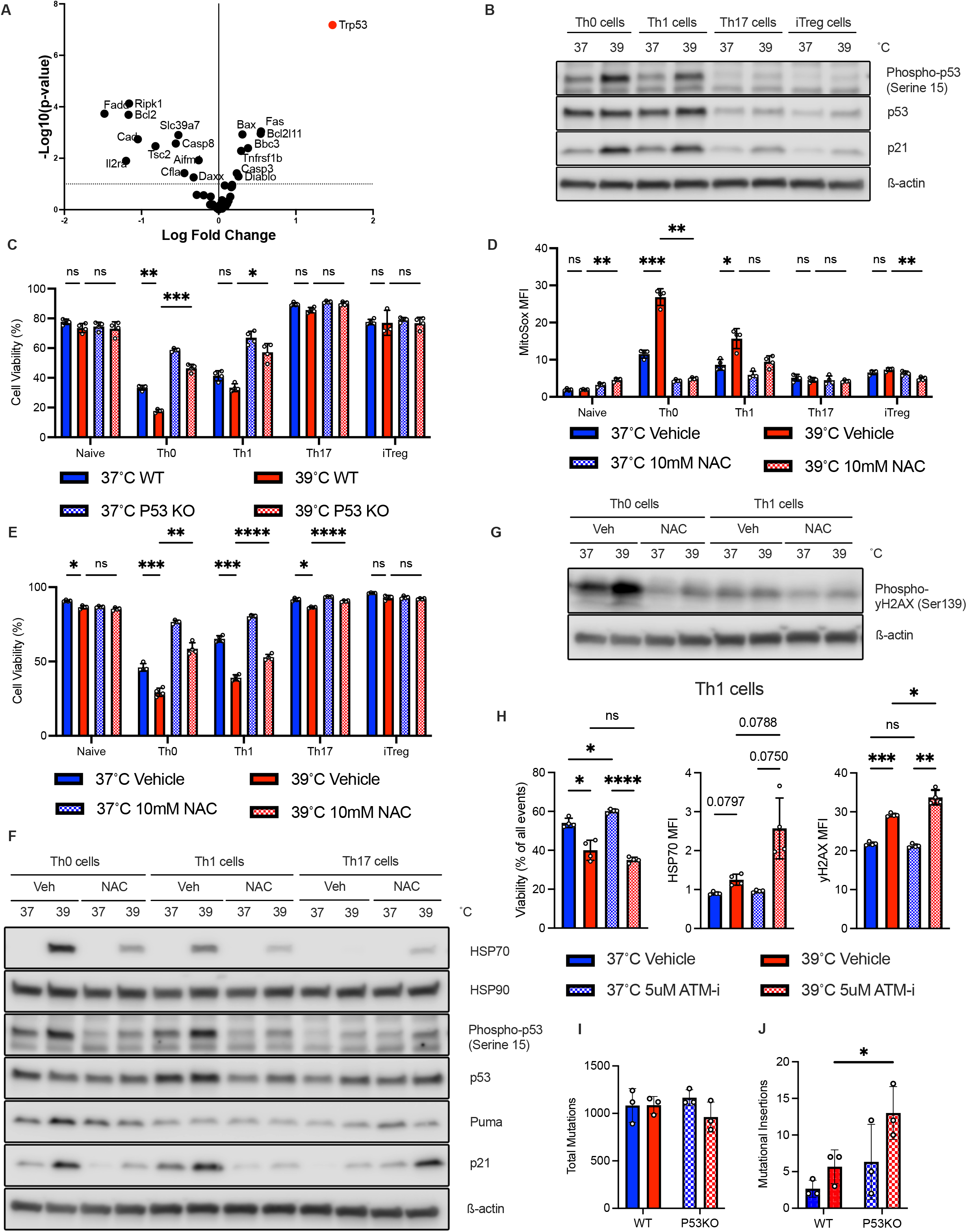
p53 is transcriptionally activated downstream of ROS induced cell stress and serves to cull mutagenic cells at 39°C. A) Cell-death CRISPR screen reveals genes providing selection advantage and disadvantage at 39°C B) Western blotting of phospho-p53-serine 15, p53, and p21 in subsets cultured at 37°C or 39°C. C) Viability of wild-type or p53 knockout cells was assessed when cultured at 37°C or 39°C D) Mitochondrial superoxide levels were measured by MitoSox flow cytometric analysis with addition or omission of ROS-Scavenger N-acetylcysteine. E) Viability of subsets cultured at 37°C or 39°C was assessed with addition or omission of NAC. F) Western blotting of previously used markers of cell stress (pP53-serine 15, HSP70, p21, Puma) with addition or omission of NAC in culture. G) γH2AX in Th0, Th1 subsets assessed by western blotting with addition or omission of NAC. H) Th1 cells were cultured +/- ATM-inhibitor at 37 or 39°C, and viability, HSP70, and γH2AX were assessed by flow cytometry. I, J) WT or p53KO Th1 cells were cultured at 37°C or 39°C for 4 days, at which point live cells were sorted and 1000X whole exome deep sequenced to identify total mutational burden (I) as well as mutational insertions (J). Biological replicates shown with standard deviation and mean. (*, P < 0.05; **, P < 0.01; ***, P < 0.001, ****, P < 0.0001; C-E: 2-Way Anova, H-J: 1-Way Anova)

To test if ROS contributed to increased cell stress, activated p53, and cell death at febrile temperatures, T cell subsets were cultured with N-acetylcysteine (NAC) to scavenge ROS. This treatment significantly decreased superoxide and general ROS (Figure 4D, S7A). Importantly, viability of Th0 and Th1 cells was significantly restored (Figure 4E) while naïve, Th17, and iTreg cells were unaffected by addition of NAC. Expression of HSP70 and phosphorylation of p53-Ser15 were also significantly reduced in Th0 and Th1 cells at 39°C when cultured with NAC (Figure 4F, S7B-D). Additionally, expression of p21 was also reduced by NAC. Consistent with ROS-driven DNA damage, γH2AX levels were reduced at 39°C in Th0 and Th1 subsets with the addition of NAC (Figure 4G, S7E). These data support a model in which heat-induced mitochondrial dysfunction and elevated ROS drive increased double stranded DNA breaks that can activate p53 that is necessary to efficiently eliminate or repair damaged Th0 and Th1 cells.

### p53 limits heat-derived mutagenesis in Th1 effector population

Given increased DNA damage and p53 activation were observed at 39°C, we hypothesized that the DNA damage response and repair would be more consequential to cells at 39°C. To test this, naïve, Th0 and Th1 cells were activated for 24 hours, then cultured with ATM inhibitor for 2 days at 37°C or 39°C. While cell viability was relatively unaffected by culture with ATM-i, HSP70 expression and γH2AX were specifically enhanced in Th0 and Th1 cells at 39°C (Figure 4H, S8A-C). Additionally, p53 was still highly phosphorylated at 39°C with ATM-i, suggesting compensatory DNA repair pathway activity (Figure S8D) and that an impaired DNA damage response is more stressful to cells at 39°C than those cultured at 37°C.

To test if increased DNA damage at 39°C required p53 to prevent accumulation of T cells with mutations, wild-type and Tp53^-/-^ T cells were subject to deep, whole exome sequencing prior to activation or after differentiation to Th1 cells at 37°C or 39°C. While total single nucleotide mutations accumulated similarly between wild-type and Tp53^-/-^ cells at both 37°C and 39°C (Figure 4I), mutational insertions that indicate double-strand DNA breaks and erroneous repair were more frequent in Tp53^-/-^ cells cultured at 39°C compared to 37°C, or to wild type T cells at either temperature (Figure 4J). These data show that T cells experience mutational stresses throughout activation and proliferation and that increased temperature drives DNA damage and mutagenesis that is mitigated at a population level by p53-dependent apoptosis or DNA repair. p53 activation thus appears critical to maintain genomic integrity of T cell populations in pro-mutagenic temperature environments.

### Human scRNA-seq analysis of inflammatory disease in vivo identifies stressed Th1-like CD4^+^ T cell population

Because heat is a cardinal feature of inflammation (*4–6*), we assessed if *in vivo* inflammation led to a similar p53-dependent DNA damage response. A published time series of gene expression analysis in mouse T cell transfer-mediated inflammatory bowel disease (IBD) (*25*) showed that heat shock protein HSP70 increased expression by the 4^th^ week of disease, with reduced expression afterwards (Figure 5A). This suggested a transient local heat stress response as inflammation progressed and began to ease. To test for a role of p53 in T cells in this *in vivo* setting, T cells were transduced with the cell death CRISPR screening library and adoptively transferred into *Rag1*^-/-^ hosts in the same model of IBD. Consistent with our *in vitro* findings, deletion of p53 increased T cell fitness and abundance in both the spleen and mesenteric lymph nodes (Figure 5B). A direct competition assay was next employed in which an equal mixture of naïve wild-type and *Tp53*^-/-^ T cells expressing CD45.1 or CD45.2, respectively, was adoptively transferred into *Rag1*^-/-^ mice to induce IBD (Figure 5C, S9A, B). As a control, Thy1.1^+^ nTregs were adoptively transferred in some *Rag1*^-/-^ mice to limit inflammation caused by transfer of a 1:1 mix of wild type and *Tp53*^-/-^ T cells. After 5 weeks, mice were sacrificed, spleen and mesenteric lymph nodes were extracted, and CD4^+^ T cells were analyzed by flow cytometry. While the ratio of wild type to *Tp53*^-/-^ cells remained unchanged in the mice receiving nTreg to prevent inflammation, *Tp53*^-/-^ T cells outcompeted WT cells in the inflammation group (Figure 5D, S9C) and this correlated with disease severity (supplemental S8D-E).

**Figure 5:**
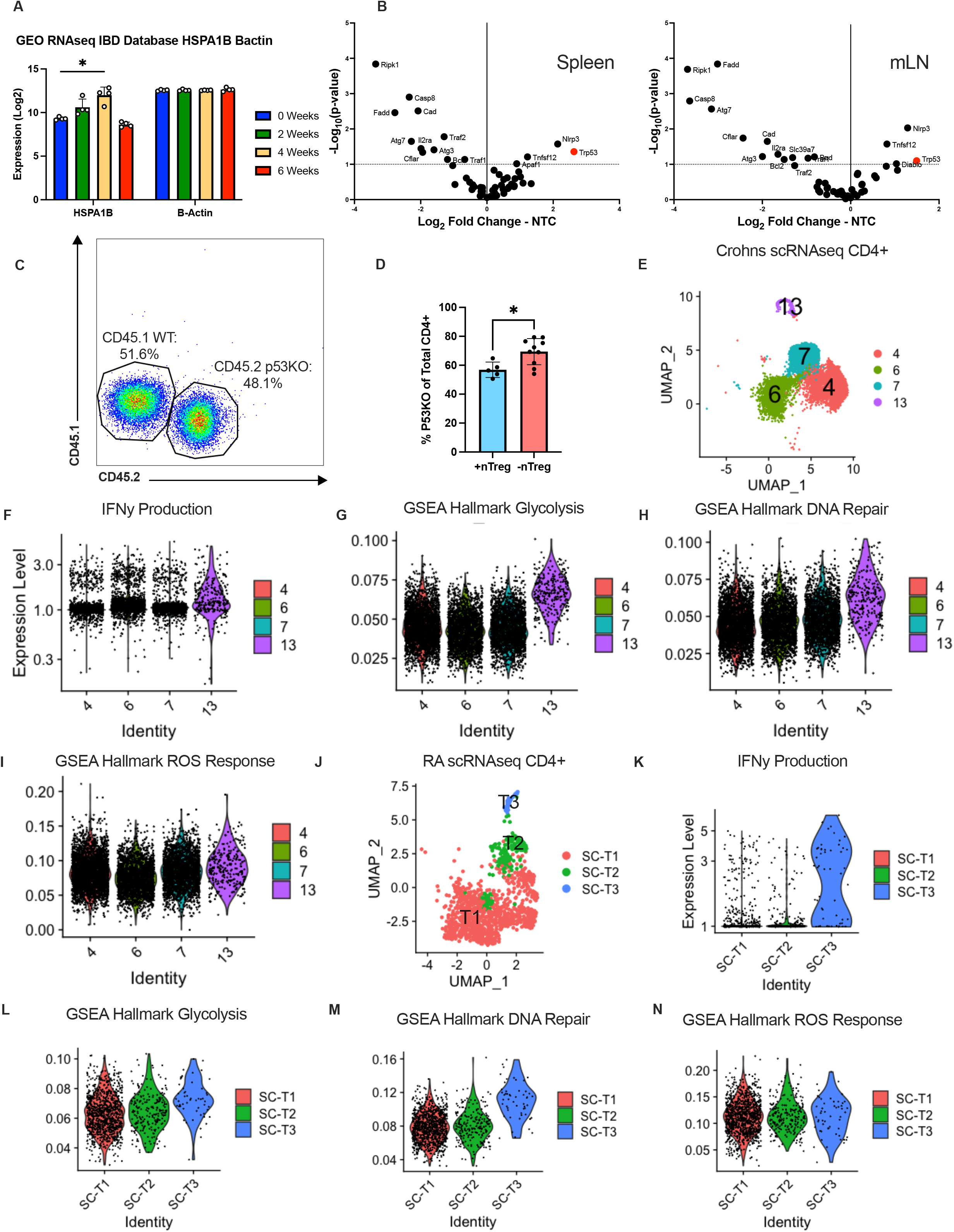
*In-vivo* correlates suggest a role for p53 and unique response to cell stress in IFNγ producing CD4^+^ T cells in inflammatory diseases associated with increased tissue temperatures. A) GEO RNAseq database was used to identify signs of heat stress response in mice with inflammatory bowel disease via HSP70 expression. B-actin used as control protein. B) In-vivo cell-death Crispr screen was applied to Th1 cells in a mouse model of IBD to identify proteins providing a survival advantage in inflammatory conditions C) A 1:1 mix of wild-type and p53KO T cells was intraperitoneally injected into RAG^-/-^ mice to test for competitive advantage in an adoptive transfer mouse model of inflammatory bowel disease. D) After 5 weeks, mice were sacrificed, and the ratio of p53KO cells to WT cells was assessed in the spleen. Percentages calculated as percentage of p53KO cells to total CD4^+^ cells. E-I) Analysis of CD4^+^ T cell groups in published human Crohn’s disease scRNAseq data J-N) Analysis of CD4^+^ T cell groups in published human rheumatoid arthritis scRNAseq data. Biological replicates shown with standard deviation and mean. (*, P < 0.05; **, P < 0.01; ***, P < 0.001, ****, P < 0.0001; A: 2-Way Anova, D: Unpaired T-Test

To identify instances of temperature change in human inflammation, we first examined published data from a scRNAseq dataset using patient samples from Crohn’s disease (*26*). We identified 4 significant CD4^+^ T cell groups from this dataset (Figure 5E, S10A, B), with cluster 4 being the greatest IFNγ producing group (Figure 5F). Like Th1 cells cultured *in vitro*, this group displayed a high gene expression signature of glycolysis (Figure 5G). Interestingly, this CD4^+^ T cell cluster also had the highest gene signature for DNA damage response (Figure 5H), one of the highest ROS responses among T cell groups (Figure 5I), as well as a high response to heat (Figure S10C). In contrast, gene set enrichment for the p53 pathway was low, which may suggest that that only IFNγ producing cells with low p53 pathway activity survive in this inflammatory setting (Figure S10D). Interestingly, inflammatory burden in inflammatory bowel disease patients has previously been identified as a risk factor for increased DNA damage and oxidative damage in T cells (*27*), which further supports a model of inflammation-induced hyperthermic stress.

To further examine *in vivo* responses to heat, we analyzed a published scRNAseq dataset from rheumatoid arthritis patients (*26*), where joints have been shown to have higher temperatures than uninflamed joints (*5, 6*). T cells in RA displayed a higher response to heat and elevated IFNγ compared to cells captured in osteoarthritis (OA) (Figure S10E, F). When CD4^+^ T cells were reanalyzed, 3 groups emerged (Figure 5J), in which IFNγ was highly expressed in cluster T3 (Figure 5K). We analyzed DNA damage repair pathway activity and found that this IFNγ producing T cell cluster again had significantly higher signature for glycolysis and response to DNA damage than the other T cell groups (Figure 5L, M). Response to ROS was also elevated in this group (Figure 5N), which again had low p53 pathway activity (Figure S10G). Taken together, these data provide correlative *in vivo* human evidence for differential response to inflammation-induced stress among CD4^+^ T cell subsets, with a population of highly glycolytic, IFNγ-producing T cells exhibiting higher DNA damage and response to ROS than other T cell groups. These data are also supportive of previous reports identifying high levels of DNA damage and apoptosis in T cells of newly diagnosed, untreated RA patients (*29–31*). While these changes cannot be directly linked to increases in temperature *in vivo*, the central role of heat in inflammation supports a model of inflammation-induced heat and DNA damage and repair in Th1 cells.

## Discussion

Temperature change is a common physiological characteristic of immune responses yet is poorly understood in the context of T cell metabolism and function. Here, we show that significant metabolic alterations and mitochondrial stress occur in T cells *in vitro* at febrile temperatures in a subset-specific manner. These differential metabolic alterations contribute to cell stress associated with increased ROS generation in Th0 and Th1 subsets, with Th17 and iTregs largely unaffected. The differential response to temperature change among T cell subsets *in vitro* raises questions as to the impact of balanced immune responses in hyperthermic settings *in vivo*. Surprisingly, increased temperatures appear mutagenic via mitochondrial stress and ROS, as heat led to p53 deficient cells exhibiting much higher levels of DNA insertions at 39°C when assessed by whole-exome sequencing. This may have many implications in cases where temperatures can increase repeatedly or for significant periods of time, including chronic inflammation, autoimmune disorders, or fevers from viral or bacterial infections. Interestingly, a recent study identified somatic mutations occurring in clonally expanded cytotoxic T cells in patients with newly diagnosed, untreated rheumatoid arthritis (*32*). The hyperthermic nature of inflammatory diseases may lead to DNA damage and somatic mutations *in-vivo* through mechanisms associated with metabolic stress and the generation of damaging reactive oxygen species.

Hyperthermia derived mutagenesis is likely not limited to rheumatoid arthritis and could occur in a variety of inflammatory contexts. High cancer burdens are common in chronic inflammation (*33*), and may be explained by cells accumulating high levels of mitochondrial stress and DNA damage (*23, 24*) while escaping a p53-mediated mechanism of cell culling. Indeed, p53 activity in T cells has been implicated in response to inflammation (*34–36*), but the underlying source of activation has not been clear. Transient heat was recently shown to lead to enhanced mitochondrial translation and a DNA damage response signature in CD8^+^ T cells (*17*), and our findings are aligned with those and suggest that longer or chronic increased temperature eventually leads to stressful ROS generation that cause DNA damage sufficient to require p53 to cull cells and maintain a less damaged pool of memory cells. While challenging to directly assess the effect of local temperature changes *in vivo* due to many off-target effects of generating temperature change, scRNAseq analysis of Th1-like CD4^+^ T cells in two different disease contexts reveal higher DNA damage response and strong similarities to data generated *in-vitro*, suggesting the presence of a heat-sensitive Th1 CD4^+^ T cells *in vivo* that correlate with our *in vitro* mechanistic studies. Further work will be needed to test for effects directly caused by temperature change *in vivo*. While this study identifies metabolic adaptations of T cells to heat, other immune cell types likely also experience metabolic perturbations at febrile temperatures. Cell-specific functional changes from heat have been well documented in literature, particularly in the innate immune system (*37*). Other cell types will be of further interest when assessing metabolic adaptations to febrile temperatures and could identify a means of targeting specific cell types in hyperthermic microenvironments.

Inflammatory diseases have long been associated with temperature change, both globally in the form of fever and locally in tissue specific context. This feature of inflammation may directly enhance the pro-inflammatory and inhibit the antiinflammatory functions of T cells. Fever can, therefore, directly support protective immunity or promote inflammatory tissue damage. Increased temperatures also lead to cell stress and DNA damage when chronic. An implication of our study is that a source of mutagenesis to initiate inflammation-associated cancers may derive from heat-dysregulated mitochondria in inflammatory environments. While DNA damage repair processes can normally repair damaged DNA, and p53 dependent apoptosis can both promote repair and cull cells with high mutational burden, some cells may escape repair or death to eventually become transformed. In this case, p53 plays a broad and frequent role in genome maintenance to mitigate inflammation-induced DNA damage.

## Acknowledgments

We would like to thank all the lab members of the J.C.R lab as well as the W.K.R. lab that assisted with this project. This work was supported in part using the resources of the Center for Innovative Technology at Vanderbilt University for mass spectrometry. TEM was performed in part with the Vanderbilt Cell Imaging Shared Resource, Vanderbilt Diabetes Research Center, Vanderbilt Digestive Disease Research Center, Vanderbilt Mouse Metabolic Phenotyping Center, and Vanderbilt Vision Center (supported by NIH grants P30CA068485, P30DK020593, P30DK058404, U2CDK059637 and P30EY008126). Genomic sequencing was performed in collaboration with the VANTAGE core services at Vanderbilt University. This work was supported through by NIH Grants R01DK105550, R01HL136664, R01CA217987, R01HL118979, and R01AI153167 to J.C.R., T32DK101003 to K.V., T32AR059039 to A.R.P., the William E. Paul Distinguished Innovator Award to JCR, and NSF GRFP award #1937963 to D.R.H.

## Author Contributions

Conceptualization: D.H, J.C.R. Genomic Sequencing: A.J., L.O, W.K., and A.G.B. Methodology: J.E., C.C., E.K., L.O., A.J., K.V., H.H., A.P, A.S., F.M., and L.B. Data analysis: X.Y, E.K., C.C., J.E., L.O., K.V., H.H., and F.M. Funding Acquisition: D.R.H, J.C.R. Supervision: J.C.R, A.G.B, C.L., and W.K.R. Writing - original draft: D.H and J.C.R. Writing - reviewing and editing: D.H., J.C.R, and J.E., K.V., L.B., and W.K.R.

## References

1. D. R. Heintzman, E. L. Fisher, J. C. Rathmell, Microenvironmental influences on T cell immunity in cancer and inflammation. Cell Mol Immunol, (2022).

2. R. D. Michalek et al., Cutting edge: distinct glycolytic and lipid oxidative metabolic programs are essential for effector and regulatory CD4+ T cell subsets. J Immunol 186, 3299–3303 (2011).

3. R. M. Daniel, M. J. Danson, R. Eisenthal, C. K. Lee, M. E. Peterson, The effect of temperature on enzyme activity: new insights and their implications. Extremophiles 12, 51–59 (2008).

4. M. Segale, The Temperature of Acutely Inflamed Peripheral Tissue. J Exp Med 29, 235–249 (1919).

5. A. Gatt et al., Thermal characteristics of rheumatoid feet in remission: Baseline data. PLoS One 15, e0243078 (2020).

6. M. Greenwald, J. Ball, K. Guerrettaz, H. Paulus, Using Dermal Temperature to Identify Rheumatoid Arthritis Patients With Radiologic Progressive Disease in Less Than One Minute. Arthritis Care Res (Hoboken) 68, 1201–1205 (2016).

7. M. D. White, C. M. Bosio, B. N. Duplantis, F. E. Nano, Human body temperature and new approaches to constructing temperature-sensitive bacterial vaccines. Cell Mol Life Sci 68, 3019–3031 (2011).

8. P. Murapa, S. Gandhapudi, H. S. Skaggs, K. D. Sarge, J. G. Woodward, Physiological fever temperature induces a protective stress response in T lymphocytes mediated by heat shock factor-1 (HSF1). J Immunol 179, 8305–8312 (2007).

9. L. Q. Gothard, M. E. Ruffner, J. G. Woodward, O. K. Park-Sarge, K. D. Sarge, Lowered temperature set point for activation of the cellular stress response in T-lymphocytes. J Biol Chem 278, 9322–9326 (2003).

10. D. Umar et al., Febrile temperature change modulates CD4 T cell differentiation via a TRPV channel-regulated Notch-dependent pathway. Proc Natl Acad Sci U S A 117, 22357–22366 (2020).

11. X. Wang et al., Febrile Temperature Critically Controls the Differentiation and Pathogenicity of T Helper 17 Cells. Immunity 52, 328–341 e325 (2020).

12. C. Lin et al., Fever Promotes T Lymphocyte Trafficking via a Thermal Sensory Pathway Involving Heat Shock Protein 90 and alpha4 Integrins. Immunity 50, 137–151 e136 (2019).

13. Q. Chen et al., Fever-range thermal stress promotes lymphocyte trafficking across high endothelial venules via an interleukin 6 trans-signaling mechanism. Nat Immunol 7, 1299–1308 (2006).

14. Q. Chen et al., Central role of IL-6 receptor signal-transducing chain gp130 in activation of L-selectin adhesion by fever-range thermal stress. Immunity 20, 59–70 (2004).

15. S. S. Evans, M. D. Bain, W. C. Wang, Fever-range hyperthermia stimulates alpha4beta7 integrin-dependent lymphocyte-endothelial adhesion. Int J Hyperthermia 16, 45–59 (2000).

16. W. C. Wang et al., Fever-range hyperthermia enhances L-selectin-dependent adhesion of lymphocytes to vascular endothelium. J Immunol 160, 961–969 (1998).

17. D. O’Sullivan et al., Fever supports CD8(+) effector T cell responses by promoting mitochondrial translation. Proc Natl Acad Sci U S A 118, (2021).

18. N. Marek-Trzonkowska et al., Mild hypothermia provides Treg stability. Sci Rep 7, 11915 (2017).

19. M. O. Johnson et al., Distinct Regulation of Th17 and Th1 Cell Differentiation by Glutaminase-Dependent Metabolism. Cell 175, 1780–1795 e1719 (2018).

20. H. P. Indo et al., A mitochondrial superoxide theory for oxidative stress diseases and aging. J Clin Biochem Nutr 56, 1–7 (2015).

21. M. S. Cooke, M. D. Evans, M. Dizdaroglu, J. Lunec, Oxidative DNA damage: mechanisms, mutation, and disease. FASEB J 17, 1195–1214 (2003).

22. M. Peluso, V. Russo, T. Mello, A. Galli, Oxidative Stress and DNA Damage in Chronic Disease and Environmental Studies. Int J Mol Sci 21, (2020).

23. S. Mateen, S. Moin, A. Q. Khan, A. Zafar, N. Fatima, Increased Reactive Oxygen Species Formation and Oxidative Stress in Rheumatoid Arthritis. PLoS One 11, e0152925 (2016).

24. O. Altindag, M. Karakoc, A. Kocyigit, H. Celik, N. Soran, Increased DNA damage and oxidative stress in patients with rheumatoid arthritis. Clin Biochem 40, 167–171 (2007).

25. K. Fang, S. Zhang, J. Glawe, M. B. Grisham, C. G. Kevil, Temporal genome expression profile analysis during t-cell-mediated colitis: identification of novel targets and pathways. Inflamm Bowel Dis 18, 1411–1423 (2012).

26. N. Jaeger et al., Single-cell analyses of Crohn’s disease tissues reveal intestinal intraepithelial T cells heterogeneity and altered subset distributions. Nat Commun 12, 1921 (2021).

27. C. Pereira et al., DNA Damage and Oxidative DNA Damage in Inflammatory Bowel Disease. J Crohns Colitis 10, 1316–1323 (2016).

28. F. Zhang et al., Defining inflammatory cell states in rheumatoid arthritis joint synovial tissues by integrating single-cell transcriptomics and mass cytometry. Nat Immunol 20, 928–942 (2019).

29. L. Shao, J. J. Goronzy, C. M. Weyand, DNA-dependent protein kinase catalytic subunit mediates T-cell loss in rheumatoid arthritis. EMBO Mol Med 2, 415–427 (2010).

30. L. Shao et al., Deficiency of the DNA repair enzyme ATM in rheumatoid arthritis. J Exp Med 206, 1435–1449 (2009).

31. Y. Li et al., Deficient Activity of the Nuclease MRE11A Induces T Cell Aging and Promotes Arthritogenic Effector Functions in Patients with Rheumatoid Arthritis. Immunity 45, 903–916 (2016).

32. P. Savola et al., Somatic mutations in clonally expanded cytotoxic T lymphocytes in patients with newly diagnosed rheumatoid arthritis. Nat Commun 8, 15869 (2017).

33. J. Kay, E. Thadhani, L. Samson, B. Engelward, Inflammation-induced DNA damage, mutations and cancer. DNA Repair (Amst) 83, 102673 (2019).

34. A. Banerjee et al., Lack of p53 Augments Antitumor Functions in Cytolytic T Cells. Cancer Res 76, 5229–5240 (2016).

35. T. M. Tran et al., A Molecular Signature in Blood Reveals a Role for p53 in Regulating Malaria-Induced Inflammation. Immunity 51, 750–765 e710 (2019).

36. S. Zhang et al., Trp53 negatively regulates autoimmunity via the STAT3-Th17 axis. FASEB J 25, 2387–2398 (2011).

37. S. S. Evans, E. A. Repasky, D. T. Fisher, Fever and the thermal regulation of immunity: the immune system feels the heat. Nat Rev Immunol 15, 335–349 (2015).

