## Supplementary material for "Subset-specific mitochondrial and DNA damage shapes T cell responses to fever and inflammation": Methods and Supplemental Figures

**Method Details**

*Mice****:*** All experiments were performed at Vanderbilt University in accordance with Institutional Animal Care and Utilization Committee (IACUC)-approved protocols and conformed to all relevant regulatory standards. Mice were housed in pathogen-free facilities in ventilated cages with ad libitum food and water and at most 5 animals per cage. Eight- to twelve-week-old male mice were used for all animal experiments. All mice were obtained from Jackson Laboratory and were treatment-naive until the start of study.

*In vitro mouse CD4^+^ T cell activation and differentiation:* Primary murine CD4*^+^* T cells were isolated from the spleens and lymph nodes of mice using a negative isolation kit according to the manufacturer’s instructions (StemCell). The cells were cultured at 37°C or 39°C with 5% CO_2_ in RPMI 1640 media (Corning) supplemented with 10mM HEPES (Corning), 50μM 2-mercaptoethanol (Sigma Aldrich), 100U/mL penicillin/streptomycin (Corning), and 2mM glutamine (Corning), unless otherwise stated. Primary CD4*^+^* T cells were activated using plate-bound anti-CD3 (3µg/mL) (Thermofisher) and anti-CD28 (5µg/mL) (Thermofisher) antibodies at 800,000-1 million cells/well for 24-well plate. Cells were cultured for 3 to 4 days as specified with subset-specific cytokines and antibodies to promote differentiation - Th1 cells: IL-12p70 (10ng/mL) (Thermofisher), IL-2 (100U/mL) (NCI) , anti-IL-4 (10µg/mL) (Bio X Cell), anti-IFNγ (1µg/mL) (Bio X Cell); Th17 cells: IL-6 (50ng/mL) (Milyteni), TGFβ (1.5ng/mL) (Peprotech), IL-23 (10ng/mL) (Milyteni), IL-1b (10ng/mL) (Milyteni), anti-IL4 (10µg/mL) (Bio X Cell), anti-IFNγ (10µg/mL) (Bio X Cell); Treg cells: TGFβ (1.5ng/mL unless otherwise noted) (Peprotech), IL-2 (100U/mL) (NCI), anti-IL4 (10µg/mL) (Bio X Cell), anti-IFNγ (10µg/mL) (Bio X Cell). CB839 (GLSi) (Sai Life Sciences LTD). was dosed at 500nM for murine cells. 2-DG (Glucose analog) (Cayman Chemical) was dosed at 2mM 3 days post-activation and incubated with cells 24 hours prior to downstream analysis. ROS scavenger rescue experiments were performed with 10mM N-acetylcysteine added to culture 24 hours post-activation.

Flow cytometry: For intracellular and transcription factor stains, cells were first stained with viability dye (eBioscience Fixable Viability Dye eFluor 780) +/- cell surface antibodies, fixed and permeabilized then stained for intracellular proteins with the appropriate kits. Transcription factor staining consisted of the eBioscience FoxP3/Transcription Factor Staining Buffer Set. For cytokine stains, cells were restimulated with 1μg/mL 12-myristate 13-acetate (PMA) (Sigma Aldrich) and 750ng/mL ionomycin (Sigma Aldrich) in the presence of GolgiPlug (Thermofisher) for four hours, then processed as other intracellular stains. Unstimulated cells served as a negative control. For mitochondrial dyes, Mitotracker Green (Thermofisher) and MitoSOX Red (Thermofisher) were combined with cell surface CD4-e450 (Thermofisher #48-0042-82) and Zombie red live/dead ef780 dye (Biolegend #65-0865-14) in complete media at 37°C with 5% CO_2_ for 25 minutes. Flow cytometry antibodies and reagents used were as follows: Cell Trace Violet (Thermofisher #C34557), Glut1 af647 (Abcam #ab195020), Mitotracker Green FM (Thermofisher #M7514), MitoSOX™ Mitochondrial Superoxide Indicator (Thermofisher #M36008), H2DCFDA (Thermofisher #D399), FITC anti-H2A.X Phospho (Ser139) (BioLegend # 613404), Alexa Fluor® 488 anti-Hsp70 Antibody (BioLegend #648004), FOXP3 Monoclonal Antibody (FJK-16s), APC (Thermofisher #17-5773-82), IFN gamma Monoclonal Antibody (4S.B3), APC (BioLegend #17-7319-41), PE anti-mouse IL-17A Antibody (BioLegend #506904), Anti-Thy1.1 ef450 (Thermofisher #48-0900-82), Anti-Brilliant Violet 510™ anti-mouse CD45.1 (BioLegend #110741), PE/Cyanine7 anti-mouse CD45.2 (BioLegend #109830). Data were analyzed using FlowJo software v10.6.2.

*iTreg Suppression Assay*: iTreg were skewed as previously described at 37°C or 39°C for 4 days. On day 2, 1ml of media was replaced with 1ml of fresh cytokine media. On day 3, iTregs were taken off CD3/CD28 stimulation plate and given 1 more mL of fresh media. Also on day 3, CD8+ T cells were isolated from WT mice and stimulated in 12-well non-tissue culture treated plates for 1 day with 100U/mL of IL-2. On day 4, CD8+ T cells were removed from plates, washed with PBS (Corning) and stained with cell trace violet (CTV) (Thermofisher) for 20 minutes at 37°C. Cells were mixed together with iTreg cells in 96-well plates (Thermofisher). No IL-2 was provided in the media. 0.5ug/ml of soluble CD28 was provided to all wells. The total cell number in all wells was 200,000 cells. iTregs were mixed with CTV effectors at ratios: 0 to 1, 1 to 4, 1 to 2, or 1 to 1 and cultured at 37°C. After 3 days, suppression assays were analyzed by flow cytometry. Cells were gated on singlets, lymphocytes, Zombie red live/dead ef780 viability dye (Thermofisher), and then either CD4-SB600 (Thermofisher) or CD8a-PE (eBioscience). CD8a positive CD4 negative cells were gated to analyze CTV proliferation peaks. All cells that divided at least once compared to the control (unstimulated CD8 T cells labeled with CTV) were quantified as divided cells for each group. The percentage of cells that had divided in each ratio was compared to the iTreg-free proliferation (0:1 ratio) sample to determine the percentage of Treg suppression.

*Transmission electron microscopy*: T cells were activated and cultured at 37°C or 39°C as described followed by transmission electron microscopy analysis through the Vanderbilt Cell Imaging Shared Resource. On day 3, cells were washed with warm PBS and then fixed in 2.5% glutaraldehyde in 0.1 M cacodylate for 1 hour at room temperature followed by 24 hours at 4°C. Following fixation, the cells were postfixed in 1% OsO4 and en bloc stained with 1% uranyl acetate the dehydrated in a graded ethanol series. Samples were gradually infiltrated with Quetol 651 based Spurr’s resin with propylene oxide as the transition solvent. The Spurr’s resin was polymerized at 60°C for 48 hours. Blocks were sectioned on a Leica UC7 ultramicrotome at 70 nm nominal thickness and the samples stained with 2% uranyl acetate and lead citrate. Transmission electron microscopy was performed using a Tecnai T12 operating at 100 kV with an AMT NanoSprint CMOS camera using AMT imaging software for single images and SerialEM for tiled datasets. Tiled datasets were reconstructed using the IMOD/eTomo software suite. Mitochondria quantification was done in FIJI by manually segmenting all mitochondria within cell cross sections from the tiled TEM datasets until at least 100 mitochondria were measured.

### Extracellular Flux Analyses: Extracellular flux analysis was performed with the Seahorse XFe96 Analyzer (Agilent). Plates were coated with Cell-Tak solution (Corning) for 30 minutes at room temperature before seeding the cells. 150,000 viable cells were seeded per well and a minimum of five technical replicates were seeded for each sample. The Mito Stress Test was performed with oligomycin A at 1.5μM, FCCP at 1.5μM, and rotenone/antimycin A at 0.5μM final concentrations.

*Lactate secretion assay* T cells were activated and cultured at 37°C or 39°C as described. 1X10^6^ cells were then seeded in 96 well plates in 200ul of fresh RPMI media and incubated for 3.5 hours at 37˚C. Media was collected for use with Abcam lactate secretion assay. The assay was performed according to manufacturer instructions. nMol Lactate/ml/hr calculation was determined by the amount of lactate present in 200ul of media after 3.5 hours, multipled by 5 and divided by 3.5 to determine nMol/ml/hr.

*CRISPR screening:* Cell death [gRNA](https://www.sciencedirect.com/topics/medicine-and-dentistry/guide-rna) library was curated by referencing the Mouse CRISPR Knockout Pooled Library (Brie) (Addgene, Pooled Library #73632). Four gRNA sequences for each gene and ten non-targeting controls, flanked by the following adaptor sequences were purchased as an oligo pool from Twist Bioscience: GGAAAGGACGAAACACCGXXXXXXXXXXXXXXXXXXXXGTTTTAGAGCTAGAAATAGCAAGTTAAAATAAGGC. The library was further prepared for transduction following published methods ([Anderson et al., 2017](https://www.sciencedirect.com/science/article/pii/S1074761321004489?via%3Dihub" \l "bib1); [Shalem et al., 2014](https://www.sciencedirect.com/science/article/pii/S1074761321004489?via%3Dihub" \l "bib40)) with several modifications. Briefly, additional sequences were attached by PCR using Array primers and Herculase II Fusion DNA Polymerase (Agilent Technologies). After purification by gel extraction using QIAquick Gel Extraction Kit (Qiagen), the fragment was cloned into the retroviral expression vector pMx-U6-gRNA-BFP using Gibson Assembly Master Mix. The resultant plasmid pool was amplified by electroporation into ElectroMAX DH10B Cells (Thermofisher) and plated on ampicillin plates to obtain enough colonies for 50-fold coverage of the library. DNA was isolated using GeneJET Plasmid Maxiprep Kit (Fisher Scientific).

DNA was transfected into Plat-E retroviral packaging cell line using Polyplus jetPRIME DNA and siRNA transfection reagent (VWR). Media was changed 24 hours post transfection, and viral supernatant was collected after an additional 48 hours of culture. Meanwhile, CD4*^+^* T cells were isolated from the spleen and lymph nodes of Cas9-transgenic mice and activated with anti-CD3 and anti-CD28. At time of CD4*^+^* T cell isolation, cells were placed in two different incubator temperatures, 37°C and 39°C. 48 hours post T cell activation, the viral supernatant was spun onto retronectin (Takara Bio) treated non-tissue culture plates at 2000xg for 2 hours at 32°C. Activated T cells were transferred to the plates and spun for an additional 20 minutes, then replaced in their respective incubators. On day 3 post T cell activation, a sample of the cells were collected at 1000-fold representation of the library from each temperature condition. Cells continued to be cultured at 37°C and 39°C until 9 days post T cell activation.

Genomic DNA from cells were extracted with Kapa Express Extract Kit (Kapa Biosystems). gRNA sequences were amplified by two rounds of PCR with two technical replicates: first round with Adaptor primers and second round with barcoded Illumina sequencing primers. The amplicons were then purified by gel extraction, combined at equimolar, and sequenced for 150 cycles in paired-end mode on the Illumina Novaseq 6000 platform at the Vanderbilt Technologies for Advanced Genomics (Vantage). At least 1000-fold representation of the library was maintained throughout the process. FASTQ files were analyzed using the Model-based Analysis of Genome-wide CRISPR/Cas9 Knockout (MAGeCK v0.5.0.3) method ([Li et al., 2014](https://www.sciencedirect.com/science/article/pii/S1074761321004489?via%3Dihub" \l "bib27)) for statistical analysis. Briefly, files are first median normalized by read count. Then, gRNA abundance across samples is compared using a negative binomial model to generate p values. Here, we compared the gRNA frequency in the CD4*^+^* T cells isolated from 39°C and 37°C at day 9 to the gRNA frequency in the CD4*^+^* T cells at day 3.

*Fluorescence Microscopy*: Cells were fixed with 4% PFA (Electron Microscopy Sciences) in PBS with 0.5% Triton-X100 (PBSTx) for 10 minutes. Cells were blocked with 1% BSA in PBSTx and stained for gH2A.X hosphor-Ser139 (1:1000, JBW301, Millipore) then goat anti-mouse AF488 (1:2000, A-11001, Invitrogen,) and DAPI. Cells were then cytospun and mounted with Prolong Gold Antifade reagent (Thermo Fisher Scientific) under #1 glass coverslips. Images were acquired using a 40 x 1.3 NA objective on a DeltaVisison Elite imaging system (GE Healthcare) equipped with a Cool SnapHQ2 charge-coupled device camera (Roper). Optical sections were collected at 200nm intervals and processed using ratio deconvolution in softWoRx (GE Healthcare). Representative images presented are maximum intensity projections, prepared in FIJI (Version 2.3.0).

*Mass spectrometry metabolomics*: Frozen samples were stored at -80°C until analyzed by LC-MS-based metabolomics. Individual samples were reconstituted in 20mL water, vortexed, and diluted with 60 mL of 90:10 acetonitrile/water containing isotopically labeled standards (carnitine-D9, valine-D8, tryptophan-D3, and inosine-4N15). Equal volumes of individual samples were pooled to create a quality control (QC) sample used for column conditioning, retention time alignment, to assess instrument reproducibility, and for batch acceptance. A Q-Exactive HF hybrid quadrupole-Orbitrap mass spectrometer (Thermo Fisher Scientific, Bremen, Germany) equipped with a Vanquish UHPLC binary system and autosampler (Thermo Fisher Scientific, Germany) was used to collect all data. Metabolite extracts were separated on ACQUITY UPLC BEH Amide HILIC column (1.7μm, 2.1 × 100 mm; Waters Corporation, Milford, MA) at 30°C. Liquid chromatography was performed using solvent A (5 mM Ammonium formate in 90% water, 10% acetonitrile and 0.1% formic acid) and solvent B (5 mM Ammonium formate in 90% acetonitrile, 10% water and 0.1% formic acid) with a 30 min gradient at 200 μL/ min. Full MS analyses were acquired over 70-1050 m/z with a 6mL injection volume (positive ion mode) or 8mL injection volume (negative ion mode), 120K resolution, a scan rate of 3.5 Hz, AGC target of 10e6, and maximum ion injection time of 100ms. MS/MS spectra were collected at 15K resolution, AGC target of 2e5, and maximum ion injection time of 100ms. Raw data were processed with Progenesis QI v.3.0 (Non-linear Dynamics, Newcastle, UK). Species were de-adducted and de-isotoped to generate unique compounds i.e., (retention time and m/z pairs). Data were normalized to all compounds.

*Immunoblotting:* Cells were lysed with base lysis buffer containing 1% IGEPAL CA-630 (Sigma Aldrich), 200mM NaCl, and 50mM Tris pH 8.0 on ice for 30 minutes. The base lysis buffer was supplemented with the protease inhibitors aprotinin (5ug/mL) (Sigma Aldrich), leupeptin (5ug/mL) (Sigma Aldrich), sodium fluoride (0.9mM) (Sigma Aldrich), dithiothreitol (DTT, 1mM) (Sigma Aldrich), sodium vanadate (1mM) (Sigma Aldrich), and beta-glycerophosphate (20mM) (Sigma Aldrich). Lysates were centrifuged for 15 minutes at 4˚C to recover supernatant and quantified for protein concentration using Protein Assay Dye Reagent Concentrate (Bio-Rad). 40ug of protein was loaded per well for polyacrylamide gel electrophoresis using Mini-PROTEAN Precast Polyacrylamide Gels (Bio-Rad). Western blotting was performed using low fluorescence PVDF membrane (Bio-Rad). Transfer was accomplished using 1X Towbin Transfer Buffer Containing 20% methanol (Thermofisher) at 300mA for 1 hour. Blots were blocked for 1 hour using 5% BSA in TBST, before incubation with primary antibody overnight at 4˚C. Blots were washed 4X times 10 minutes each wash with TBST. Blots were incubated in 5% milk containing anti-rabbit or anti-mouse HRP-linked secondary antibody (Cell Signaling Technologies) for 30 minutes. Blots were washed in TBST, incubated with Pierce ECL western blotting substrate (Thermofisher), then imaged on a GE Healthcare Amersham Imager 600 series. The antibodies used for westerns were: ß-actin (1:1000) (CST #4967), phosphor-S6 (Ser235/236) (1:2000) (CST #4858), S6 (1:1000) (CST #2317), phosphor-Akt(Ser473) (1:1000) (CST #9271), Akt (1:1000) (CST #2920), phosphor-4EBP1(Thr37/46) (1:1000) (CST #2855), 4EBP1 (1:1000) (CST #9644), HSP70 (1:1000) (Enzo Life Sciences), HSP90 (1:1000) (CST #4874), yH2AX (Ser139) (1:1000) (CST #9718), Puma (1:1000) (CST #4976), p21 (1:1000) (Abcam AB188224), phosphor-p53 (ser15) (1:1000) (CST #9284), p53 (1C12) (1:1000) (CST #2524).

*In vivo IBD models for competition analysis of P53KO T cells vs. wild type:* Donor cells were prepared by isolating naïve CD4*^+^* T cells (CD4^+^CD25^-^CD45RB^hi^) by magnetic bead isolation (Miltenyi) from C57BL/6 WT and *Tp53-/-* (Jackson Labs; P53KO) mice aged 8-12 weeks. Thy1.1+ natural Tregs were prepared by isolating CD4+ nTregs using the CD25+ Treg isolation kit from Milyteni. CD45.1 WT and CD45.2 P53KO cells were mixed 1:1 in PBS and split into 2 tubes at 1.6X10^6^/ml final concentration. To control for inflammation induced competition, natural Tregs were added to the control mix tube at 2X10^6^ final concentration. 200,000 CD45.1 WT and CD45.2 P53KO cells each were were injected into the peritoneum of 8-week old Rag^-/-^ mice, in addition to 500,000 nTregs in the control group. Rag^-/-^ mice were housed for 5 weeks post-injection. Mice were weighed twice per week to monitor disease. Mice were euthanized after 5 weeks and spleen, colon, and mesenteric lymph nodes were collected for T cell phenotyping by flow cytometry.

*1000X Deep Sequencing and Mutations Analysis*: Upon completion of sequencing, FASTQ files were automatically generated from bcl files upon the completion of the NovaSeq 6000 run using BaseSpace’s FASTQ Generation | Version: 1.0.0. Prior to running analysis, the custom reference, GCF_000001635.27_GRCm39 genomic.fa was imported using BaseSpace’s FASTA Upload | Version: 1.0.0.  In order to upload the reference, the file extension was updated from .fna to .fa. This reference is available for download via ([https://www.ncbi.nlm.nih.gov/assembly/GCF_000001635.27/](https://nam04.safelinks.protection.outlook.com/?url=https%3A%2F%2Fwww.ncbi.nlm.nih.gov%2Fassembly%2FGCF_000001635.27%2F&data=05%7C01%7Cdarren.r.heintzman%40vanderbilt.edu%7C909eb74533b141d605d008dabea12c32%7Cba5a7f39e3be4ab3b45067fa80faecad%7C0%7C0%7C638031896693688955%7CUnknown%7CTWFpbGZsb3d8eyJWIjoiMC4wLjAwMDAiLCJQIjoiV2luMzIiLCJBTiI6Ik1haWwiLCJXVCI6Mn0%3D%7C3000%7C%7C%7C&sdata=FAPNnnG%2Bh2WMUeuLUJnygpi1VymIZUU%2FynC7aSambkA%3D&reserved=0)). Once the reference was upload, BaseSpace DRAGEN Reference Builder | Version: 3.10.4 was used to build the required V8 Hash Table that Illumina BaseSpace DRAGEN applications require for downstream analysis. Genomic sequences of cells cultured at 37˚C or 39˚C were compared to day 0 isolated naïve cells from the same biological replicate. For analysis, Illumina’s BaseSpace DRAGEN Somatic | Version: 4.0.3 with the following parameters was used to complete the required comparisons for this manuscript: under Advanced Settings, Somatic Hotspots: None (since an imported/custom reference was used), and Under Additional Arguments, check the acknowledgement statement and add the following command, --vc-enable-unequal-ntd-errors=false (required per discussion with Illumina Tech-Support, recommended due to software memory limitations.  If not included, analysis may abort unexpectedly.)

*scRNAseq of Crohns and RA data*: The Crohn’s disease single-cell RNAseq data gene expression matrix was downloaded from GEO under the GSE157477 (*1*). Aggregated gene expression matrices containing numbers of UMIs per cell per gene were filtered to retain cells with at least 500 genes detected and less than 15% of total UMIs originating from mitochondrial RNA. Genes detected in more than 3 cells were retained for the following analysis. The Rheumatoid arthritis single-cell RNAseq data gene expression matrix was downloaded from ImmPort (https://www.immport.org/shared/study/SDY998, study accession code SDY998) (*2*). Then the gene expression matrix was filtered to retain genes expressed in at least 10 cells and cells with at least 1000 genes detected and less than 25% of total mitochondrial RNA. Dimension reduction (principal component analysis [PCA], UMAP) and clustering were applied to the filtered matrix using Seurat (v4.0.6) with default parameters, except the top 20 dimensions of PCA were used for UMAP. R package AUCell (v1.16.0) was used to calculate AUC scores of HALLMARK and GO pathways for each cell.

**Quantification and Statistical Analysis**

Statistical analyses were performed with Prism software (v8). Statistically significant results are labelled (* p < 0.05, ** p < 0.01, *** p < 0.001, **** p ≤ 0.001). Error bars show mean ± standard deviation unless otherwise indicated. All experiments were carried out in quadruplicate biological replicates unless otherwise stated. Flow cytometric plots shown are representative of biological replicates.

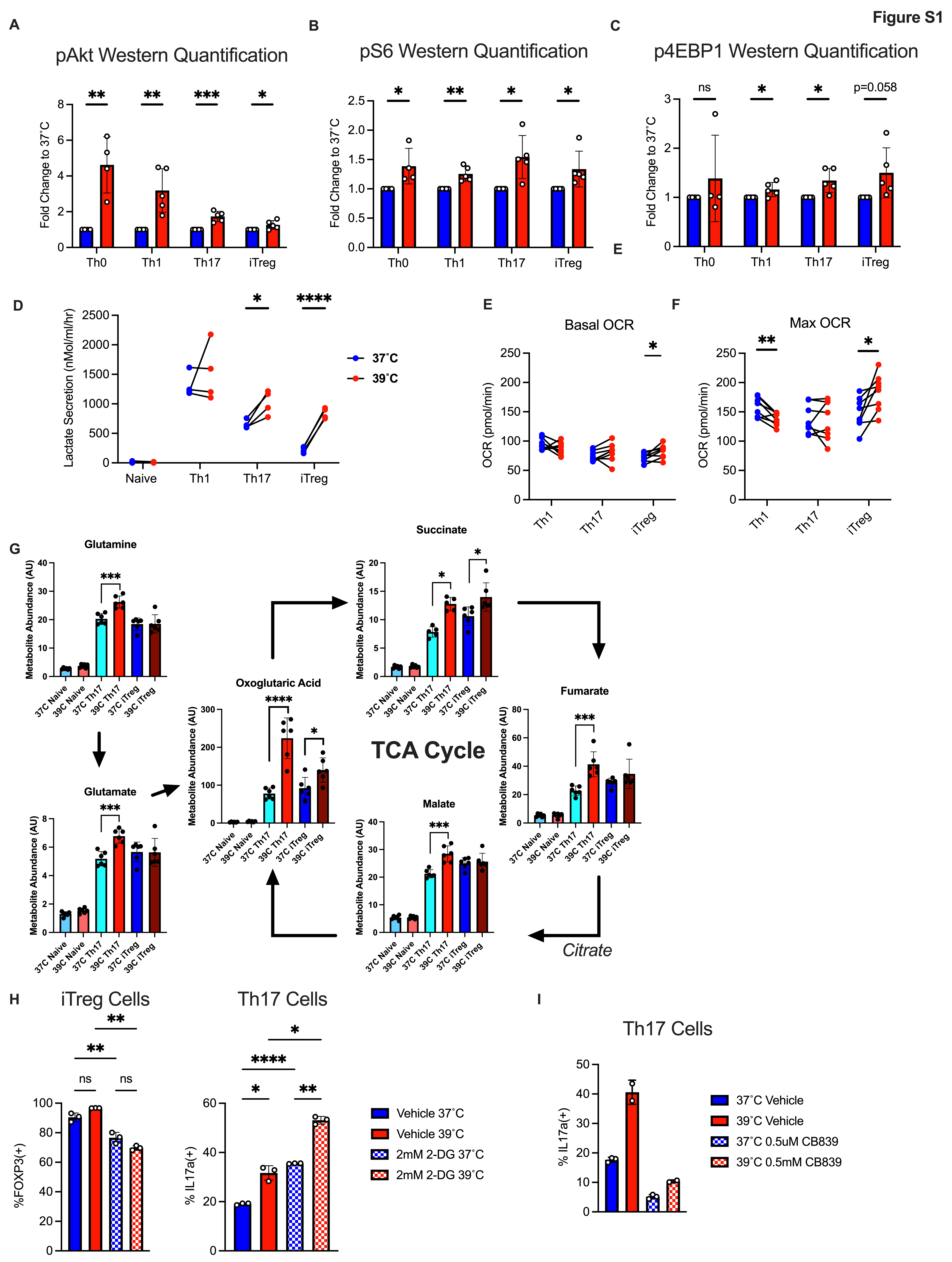

**Figure S1**: **CD4 T cell metabolism is differentially enhanced at febrile temperatures**. A-C) Quantification of mTor pathway proteins from western blot shown in Figure 2A. D) Lactate secretion in naïve, Th1, Th17, and iTreg cells cultured at 37˚C and 39˚C. E,F) Basal and maximal oxidative consumption rates measured by Seahorse. G) Percentage of FOXP3(+) cells in iTregs cultured at 37 or 39˚C with or without 2-DG. H) Percentage of IL17a(+) cells in Th17 cultures with or without 2-DG at 37 or 39˚C. G) Percentage of IL17a(+) Th17 cells cultured with or without hTGF-b at 37 or 39˚C. I) Targeted metabolomics was performed on naïve, Th17, and iTreg cells at 37 and 39˚C for glutaminolysis and TCA cycle metabolites. J) IL17a expression in Th17 cells cultured with or without CB839 at 37˚C or 39˚C. Biological replicates shown with standard deviation and mean. (*, P < 0.05; **, P < 0.01; ***, P < 0.001, ****, P < 0.0001;, A-C, H: Unpaired T-Test D, I-J: Paired T-Test E, F: 1-Way Anova)

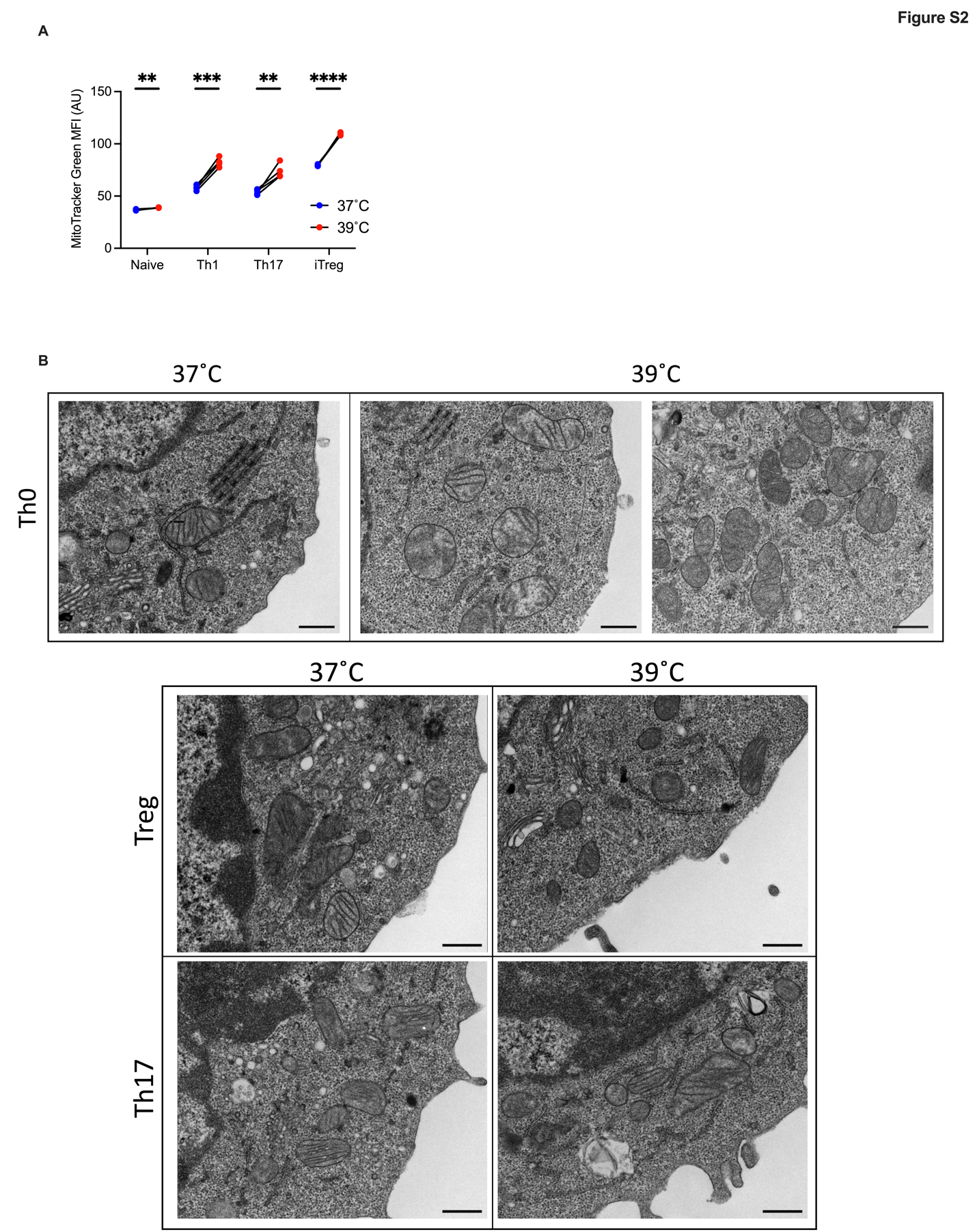

**Figure S2**: **Electron Microscopy reveals differential mitochondrial morphology in CD4+ T cells at 39˚C**. A ) Mitochondrial mass measured using MitoTracker green by flow cytometry. B) Electron microscopy images of Th0, iTreg, and Th17 cell mitochondria at 37˚C and 39˚C. Biological replicates shown with standard deviation and mean. (*, P < 0.05; **, P < 0.01; ***, P < 0.001, ****, P < 0.0001; A: Paired T-Test)

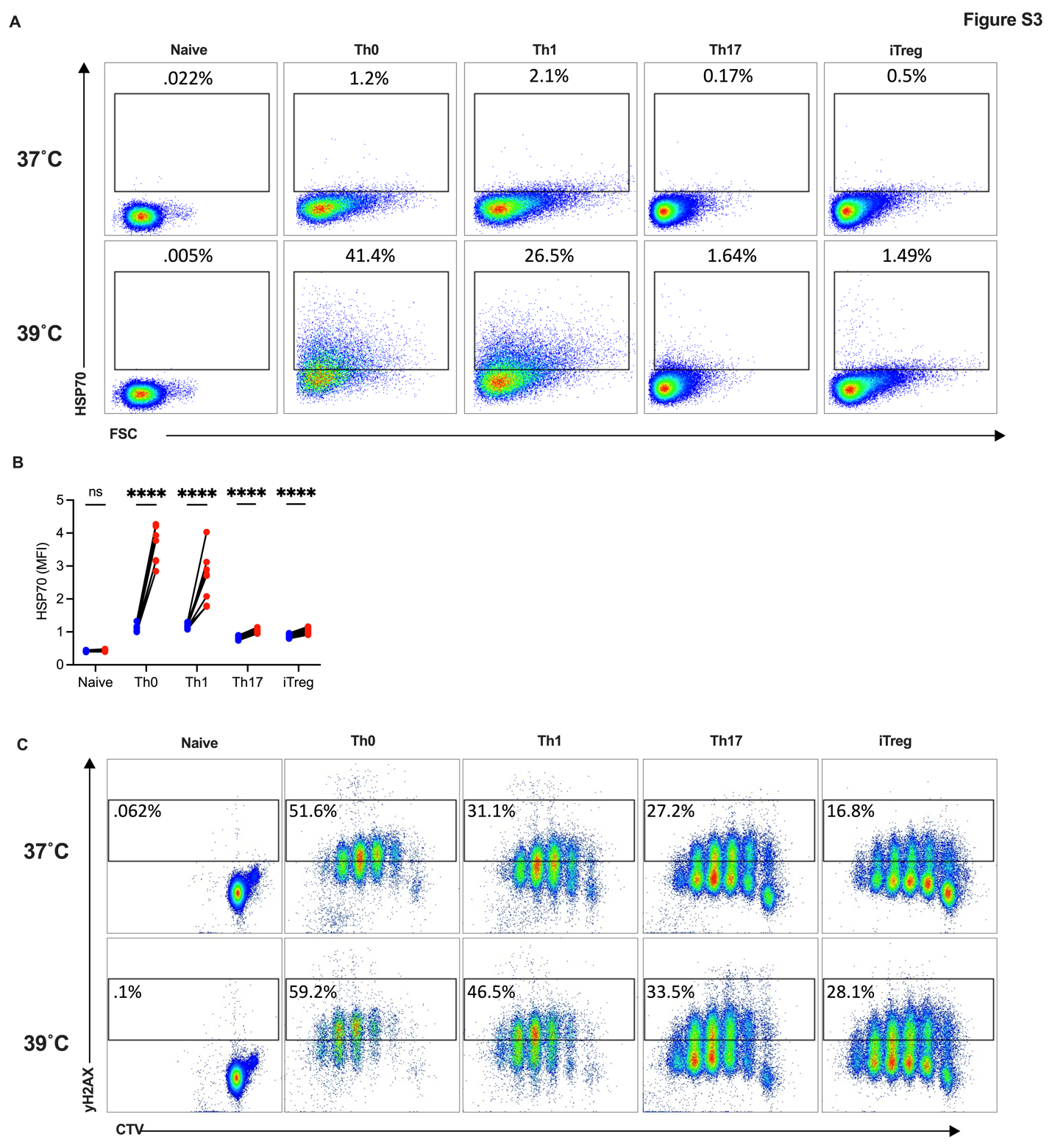

**Figure S3**: **Cell stress is markedly higher in Th0 and Th1 cells at 39˚C**. A) Flow cytometry plots of HSP70 in naïve, Th0, Th1, Th17, and iTreg cells at 37˚C and 39˚C. Plots gated on live, CD4+ cells. B) Flow cytometry quantification of HSP70 MFI in all subsets C) Flow cytometry plots of yH2AX plotted against CTV staining to identify yH2AX intensity in each cell division. Biological replicates shown with standard deviation and mean. (*, P < 0.05; **, P < 0.01; ***, P < 0.001, ****, P < 0.0001; B: Paired T-Test)

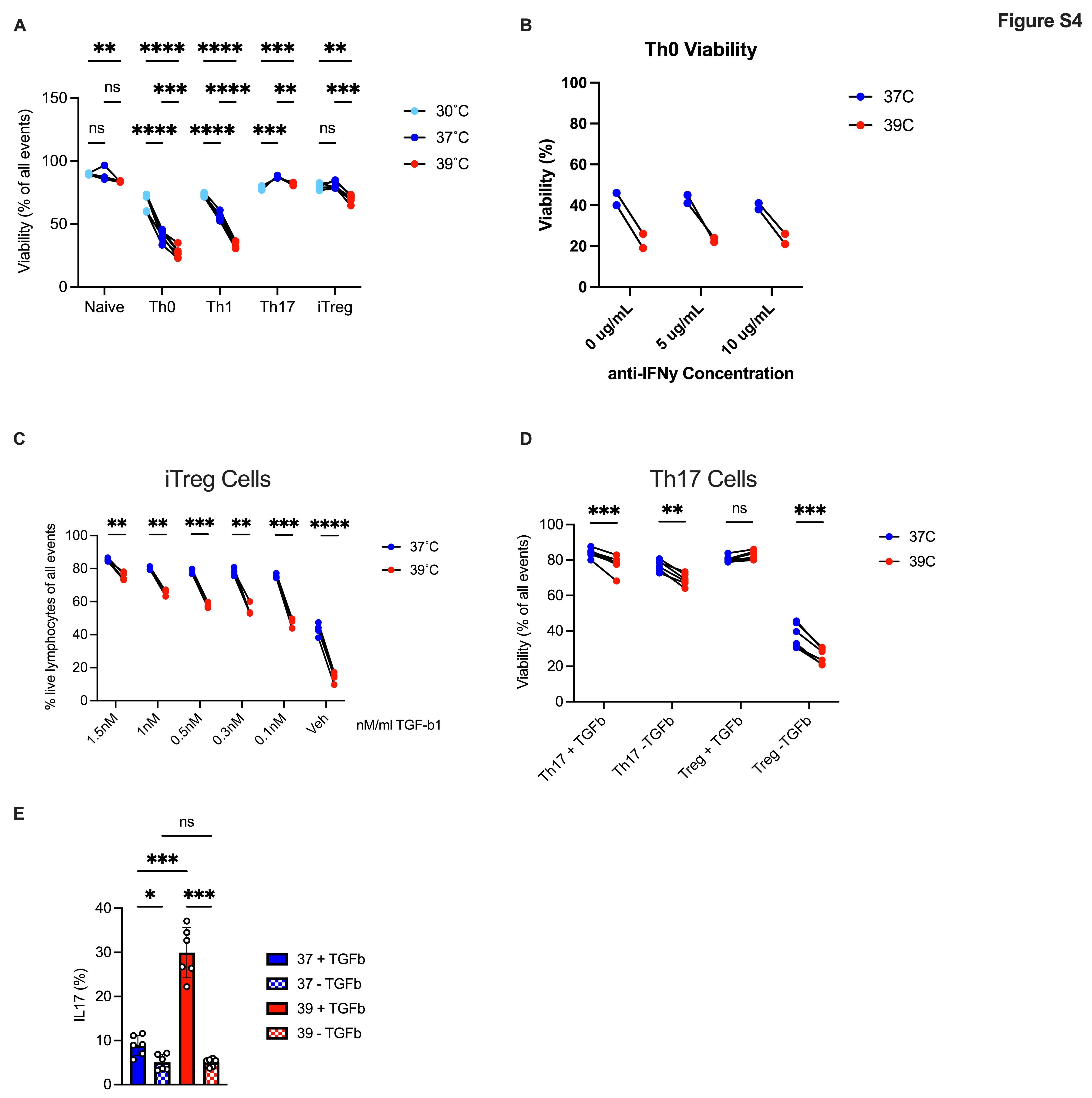

**Figure S4**: **Th0 and Th1 cell viability is specifically reduced at febrile temperatures** A) Viability of CD4+ T cells cultured at 30, 37, and 39˚C. B) Viabiilty of Th0 cells cultured with increasing concentrations of anti-IFNy antibody. C) iTreg cell viability in cultures titrated with differing amounts of hTGF-b. D) Th17 and iTreg viability was measured with or without TGF-b included in culture. E) Percentage of IL17a(+) Th17 cells cultured with or without hTGFb at 37 or 39˚C. Biological replicates shown with standard deviation and mean. (*, P < 0.05; **, P < 0.01; ***, P < 0.001, ****, P < 0.0001; A-D: Paired T-Test, E: 1-Way Anova)

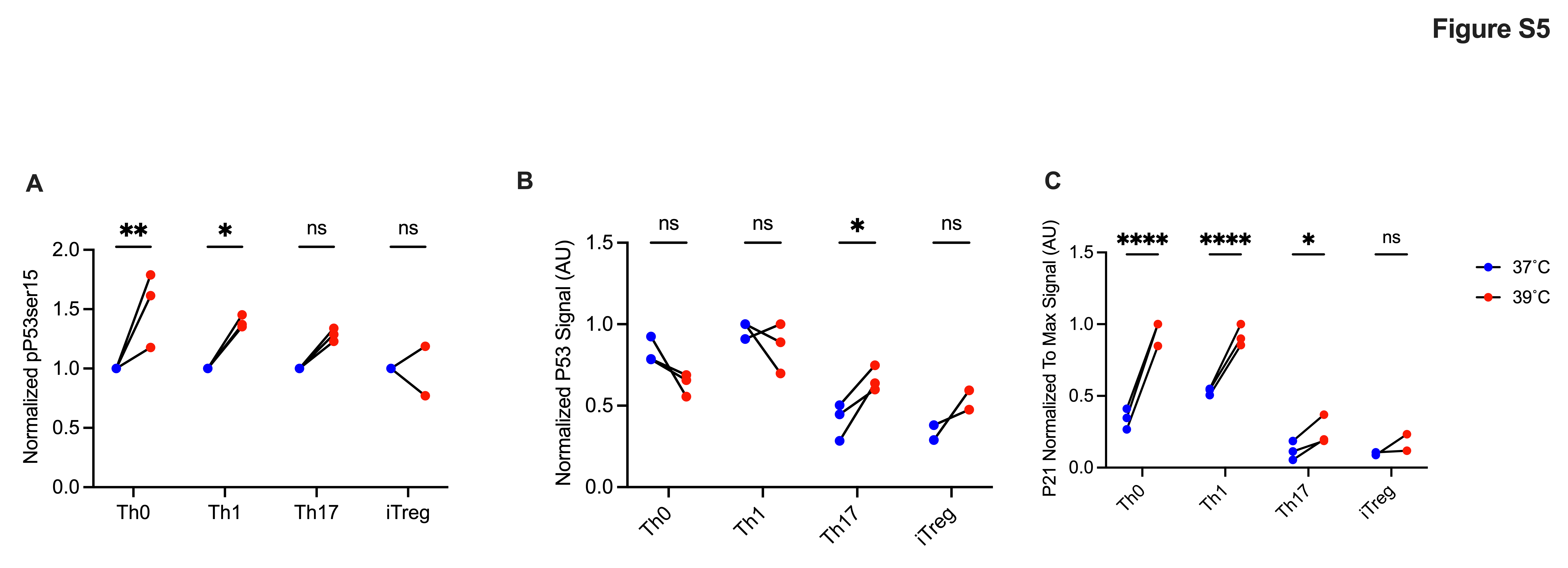

**Figure S5: p53 pathway activation occurs specifically in Th0 and Th1 subsets at febrile temperatures** A-C) Western blot quantification from Figure 3B of phosphor-P53(ser15), total p53, and p21 proteins in Th0, Th1, Th17, and iTreg cells cultured at 37 or 39˚C. Biological replicates shown with standard deviation and mean. (*, P < 0.05; **, P < 0.01; ***, P < 0.001, ****, P < 0.0001; A-C: Paired T-Test)

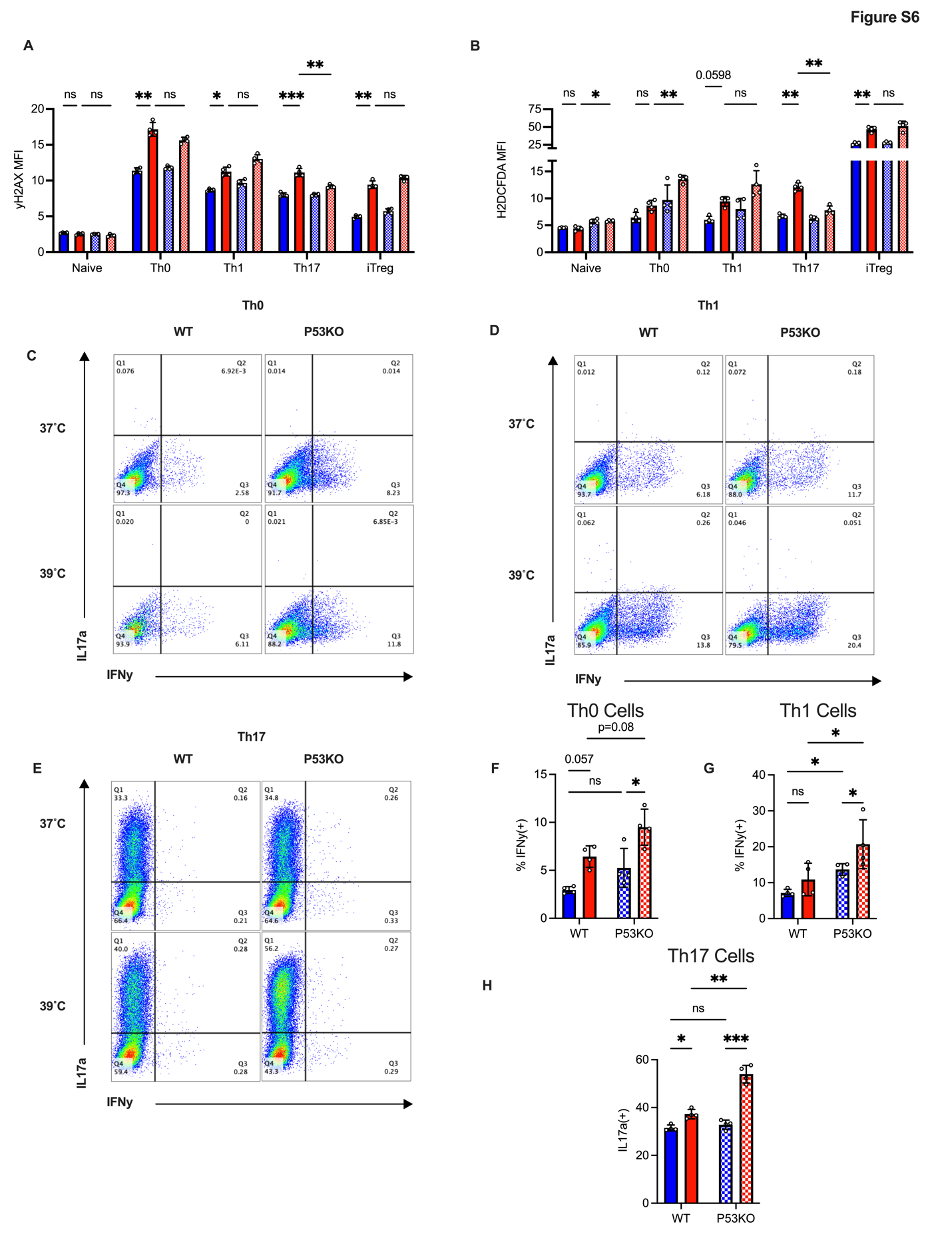

**Figure S6**: **p53 knockout leads to equally stressed cells at 39˚C but better effector characteristics**. A) yH2AX MFI quantification in all subsets, wild-type or p53KO, at 37˚C or 39˚C B) H2DCFDA general ROS staining in all subsets, wild-type or p53KO, at 37˚C or 39˚C C) IFNy production in WT vs p53KO Th0 cells at 37˚C or 39˚C D) IFNy production in WT vs p53KO Th1 cells at 37˚C or 39˚C E) IL17a production in WT vs p53KO Th17 cells at 37˚C or 39˚C F) Quantification of C G) Quantification of D H) Quantification of (E). Biological replicates shown with standard deviation and mean. (*, P < 0.05; **, P < 0.01; ***, P < 0.001, ****, P < 0.0001; A-B: 2-Way Anova, F-H: 1-Way Anova)

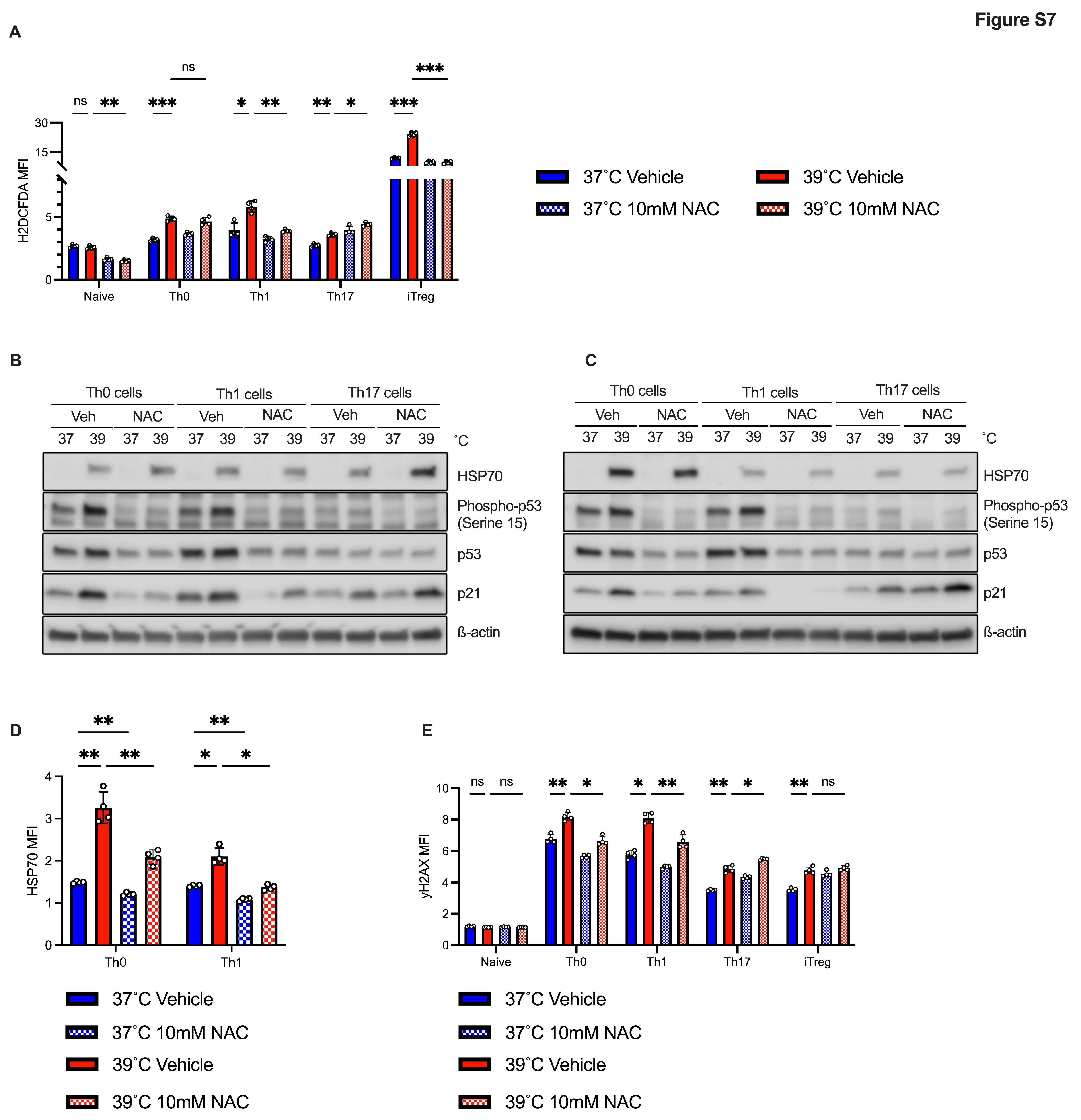

**Figure S7**: **ROS-mediated** **DNA damage is elevated in Th0, Th1 cells at 39˚C and contributes to DNA damage response** A) General cytoplasmic ROS assessed by flow cytometry of T cells cultured with or without N-Acetylcysteine (NAC). B,C) Replicate western blots of NAC experiment. D) Flow cytometry quantification of HSP70 expression in Th0 and Th1 cells cultured with or without NAC. E) yH2AX measured by flow cytometry in cells cultured with or without NAC at 37 or 39˚C. Biological replicates shown with standard deviation and mean. (*, P < 0.05; **, P < 0.01; ***, P < 0.001, ****, P < 0.0001; A,D-E: 2-Way ANOVA).

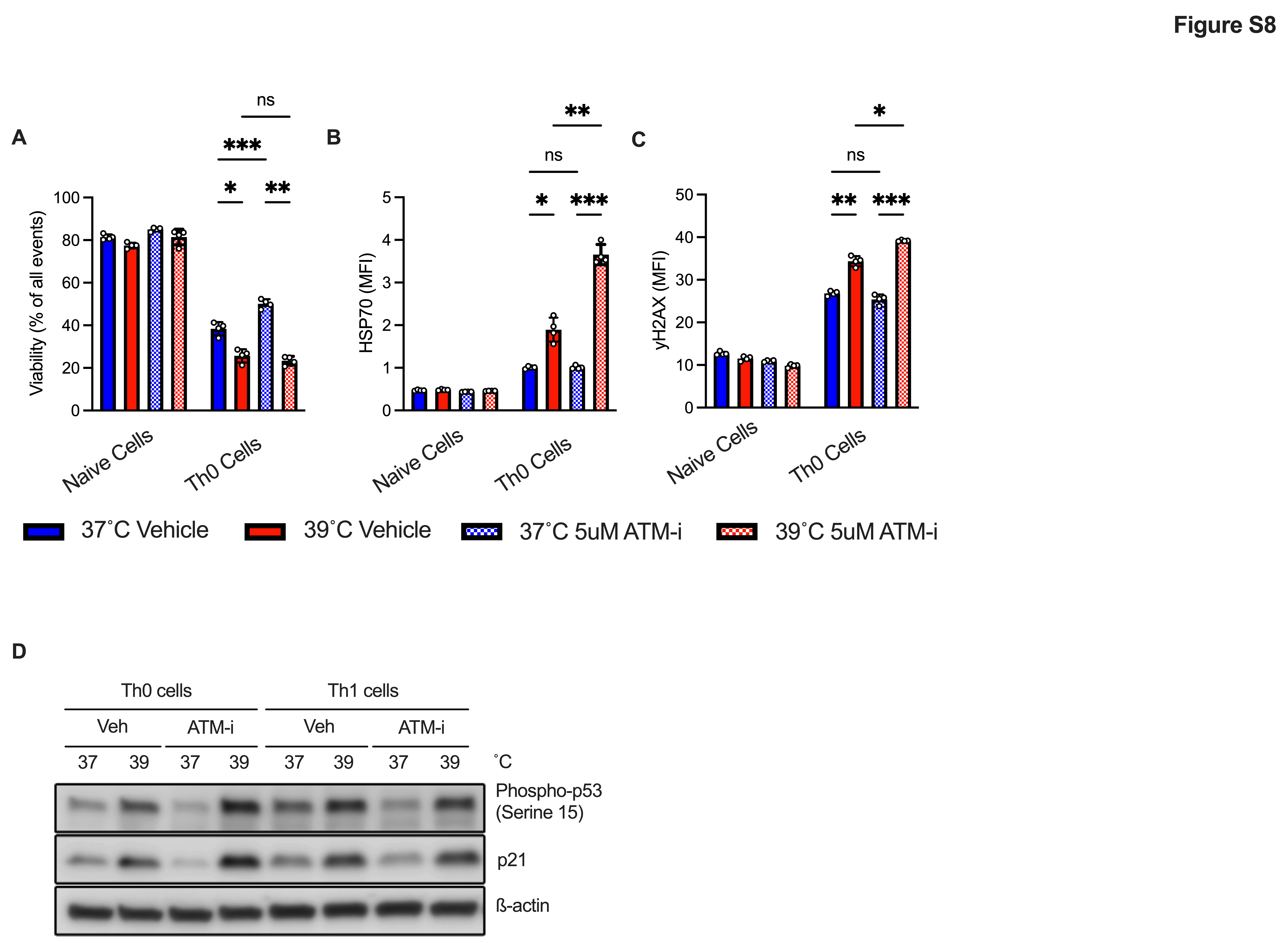

**Figure S8: DNA damage response is highly active in Th0 and Th1 subsets at febrile temperatures** A) Viability of naïve, Th0 cells after culture with or without ATM-I at 37 or 39˚C. B) HSP70 expression (MFI) quantified by flow cytometry in naïve, Th0 cells cultured with or without ATM-i at 37 or 39˚C. C) yH2AX MFI quantified by flow cytometry of naïve, Th0 cells cultured with or without ATM-i at 37 or 39˚C. D) Western blot of phosphor-p53(ser15) and p21 in Th0, Th1 cells cultured with or without ATM-i at 37 or 39˚C. Biological replicates shown with standard deviation and mean. (*, P < 0.05; **, P < 0.01; ***, P < 0.001, ****, P < 0.0001; A-C: 2-Way ANOVA,

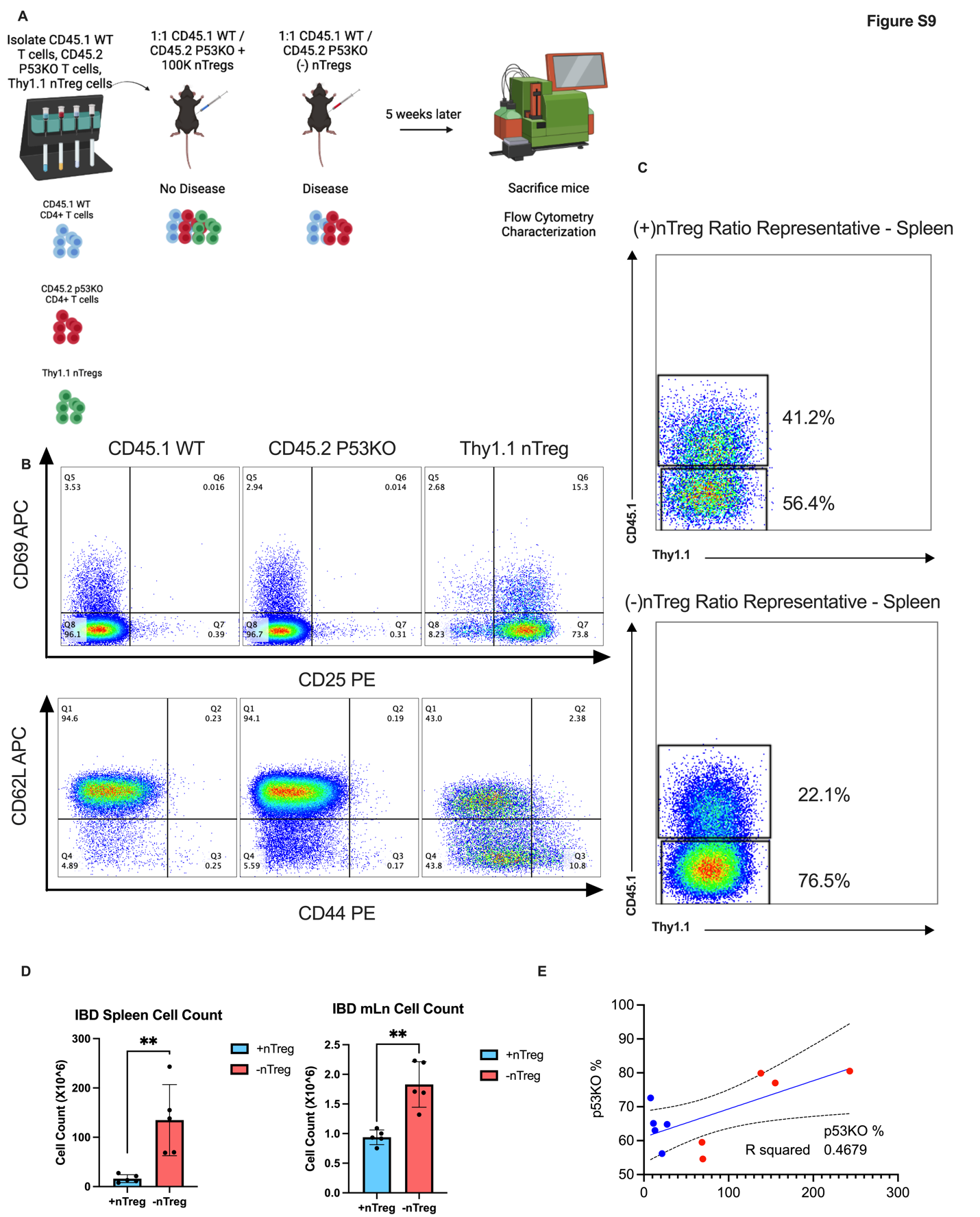

**Figure S9:** **p53 Knockout CD4+ T Cells Outcompete WT cells in a mouse model of inflammatory bowel disease.** A) Experimental design of 1:1 WT, p53KO intraperitoneal injection experiment. B) Flow cytometry plots showing activation markers of naïve cells isolated from WT, p53KO mice, and nTreg cells isolated from Thy1.1 mice. C) Representative flow cytometry plots of WT:p53 ratio analysis in spleen. nTregs were gated out of the representative control plot. D) Cell counts were quantified using trypan blue after dissection of spleen and mesenteric lymph node. E) p53KO cell ratio was compared to total cell counts of spleen within each mouse to identify a relationship between disease severity and p53KO out competition. Biological replicates shown with standard deviation and mean. (*, P < 0.05; **, P < 0.01; ***, P < 0.001, ****, P < 0.0001; D: Unpaired T-Test.

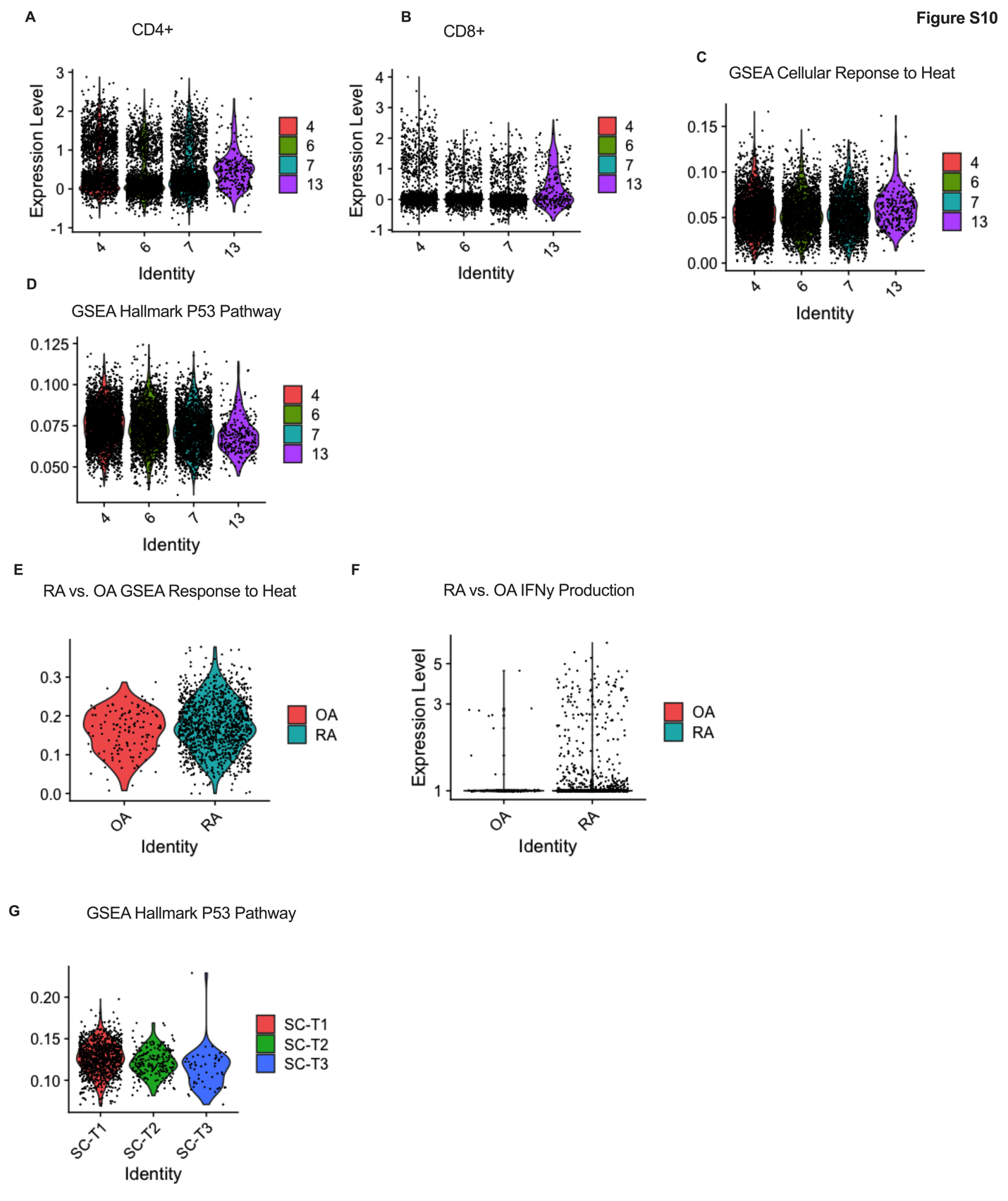

**Figure S10**: ***in-vivo* scRNASeq datasets identify an IFNy producing CD4 T cell group that displays heat stress and correlates with cells cultured *in-vitro*** A) CD4+ T cell groups were reanalyzed from the Crohn’s disease dataset. B) CD8+ T cells were reanalyzed within the CD4+ groups of the Crohn’s disease dataset. C) GSEA cellular response to heat was analyzed within the CD4+ T cell groups of the Crohn’s disease data D) p53 Pathway activity in CD4+ T cell groups of the Crohn’s disease dataset E) GSEA cellular response to heat in cells of osteoarthritis control scRNAseq and rheumatoid arthritis cells from scRNAseq dataset. F) GSEA IFNy expression in cells of osteoarthritis control scRNAseq and rheumatoid arthritis cells from scRNAseq dataset G) GSEA hallmark p53 activity in CD4+ T cell groups of rheumatoid arthritis scRNAseq dataset.

**Supplementary References**

1.	N. Jaeger *et al.*, Single-cell analyses of Crohn's disease tissues reveal intestinal intraepithelial T cells heterogeneity and altered subset distributions. *Nat Commun* **12**, 1921 (2021).

2.	F. Zhang *et al.*, Defining inflammatory cell states in rheumatoid arthritis joint synovial tissues by integrating single-cell transcriptomics and mass cytometry. *Nat Immunol* **20**, 928-942 (2019).
